# Emergence of novel non-aggregative variants under negative frequency-dependent selection in *Klebsiella variicola*

**DOI:** 10.1101/2023.07.10.548335

**Authors:** Amandine Nucci, Juliette Janaszkiewicz, Eduardo P.C. Rocha, Olaya Rendueles

## Abstract

*Klebsiella variicola* is an emergent human pathogen causing diverse infections, including in the urinary tract. However, little is known about the evolution and maintenance of genetic diversity in this species, the molecular mechanisms and their population dynamics. Here, we characterized the emergence of a novel rdar-like morphotype which is contingent both on the genetic background and the environment. We show that mutations in either the nitrogen assimilation control gene *(nac)* or the type III fimbriae regulator, *mrkH,* suffice to generate rdar-like colonies. These morphotypes are primarily selected for the reduced inter-cellular aggregation as a result of loss-of-function yielding reduced fimbriae expression. Additionally, these clones also display increased growth rate and reduced biofilm formation. Direct competitions between rdar and wild type clone show that mutations in *mrkH* provide large fitness advantages. In artificial urine, the morphotype is under strong negative frequency-dependent selection and is able to socially exploit wild type strains. An exhaustive search for *mrkH* mutants in public databases revealed that *ca* 8% of natural isolates analysed had truncated MrkH proteins many of which were due to insertions of IS elements, including a reported clinical isolate with rdar morphology. These strains were all isolated from human, mostly from urine. The decreased aggregation of these mutants could have important clinical implications as such clones could better disperse within the host allowing colonisation of other body sites and leading to systemic infections.

**One-sentence Summary:** Report of the emergence of a novel non-aggregative colony morphology in *K. variicola* and the first example of social exploitation in the *Klebsiella* genus.

## INTRODUCTION

How diversity emerges and is maintained in microbial populations are central questions in evolution and ecology. During evolution in sympatry, diversity can be driven by genetic drift, epistasis, or intercellular interactions (mostly competition) (Kassen, 2014). For instance, competition for resources can select for key innovations allowing the use of previously unavailable nutrients (Blount, Borland and Lenski, 2008). Similarly, historical contingency, that is, existing mutations in one clone may alter the range of subsequent mutations available and defining different adaptive outcomes (Blount, Borland and Lenski, 2008; Blount *et al*., 2012; Debray, De Luna and Koskella, 2022; Batarseh *et al*., 2023). One of the most potent forces of diversification is divergent selection. This is exemplified by structured environments where microniches can be generated owing to gradients of nutrient or oxygen (Rainey and Travisano, 1998). Thus, adaptive morphotypes may emerge in different microniches giving rise to specialists (Kassen, 2002; Baquero *et al*., 2021). Additionally, the existence of microniches may support growth of subpopulations with lower effective sizes which may be more subject to genetic drift. Similarly, balancing selection, by which polymorphism in a given locus is sustained, can promote genetic diversification (Hedrick, 2007). This is specially so under temporally or spatially variable environments or in fluctuating environments (Abdul-Rahman, Tranchina and Gresham, 2021).

Once it has emerged, microbial diversity can be maintained by the aforementioned spatial structure (Comins and Hassell, 1996; Kassen and Rainey, 2004) that reduces or limits migration across subpopulations. This allows for the coexistence of different morphotypes even when this could not be so in well-mixed environments (Habets *et al*., 2006). Diversity can also be maintained by cross-feeding (D’Souza *et al*., 2018), interference competition (Czárán, Hoekstra and Pagie, 2002; Kerr *et al*., 2002), dormancy (Jones and Lennon, 2010), positive (Rendueles, Amherd and Velicer, 2015) and negative frequency-dependent selection (Rainey *et al*., 2000; Velicer, Kroos and Lenski, 2000; Lemonnier *et al*., 2008). Specifically, positive-frequency dependent selection contributes to global diversity allowing genotypes that are less fit when present at intermediate frequencies to persist in patchily distributed populations provided that they are locally more abundant (Rendueles, Amherd and Velicer, 2015). Negative frequency-dependent selection occurs when increasing abundance results in decreasing fitness. This allows relatively rare variants to be maintained because they have a selective advantage over more common variants, avoiding local extinction.

For over forty years, evolution experiments have fueled the phenotypic and subsequent genotypic studies of bacterial diversification. Trait diversification has been documented in many of the phenotypes studied including cell size (Travisano, Vasi and Lenski, 1995; Baselga-Cervera *et al*., 2023), resource utilization (Tyerman *et al*., 2005; Spencer *et al*., 2008) or swarming rate (Rendueles and Velicer, 2017), among others (Travisano, 1997; Rendueles and Velicer, 2020; La Fortezza *et al*., 2022). One of the most visible features of these evolution experiments is the emergence of different colony morphotypes. Among the best well-known are the so-called wrinkly (WS) and fuzzy spreaders (FS) in *Pseudomonas flourescens* when grown in static microcosms of nutrient-rich medium (Rainey and Travisano, 1998). Similar wrinkly or rough phenotypes also emerge during growth of *Burkholderia cenocepacia* as a biofilm (Poltak and Cooper, 2011) or of *Bacillus subtilis* cells during colonization of *Arabidopsis thaliana* roots (Blake *et al*., 2021). Such rapid diversification also occurs in more natural settings complex soil microcosms (Gómez and Buckling, 2013) or as exemplified by the variety of *E. coli* colonies varying in size and motility during gut colonization (De Paepe *et al*., 2011).

The *Klebsiella pneumoniae* species complex is a metabolically versatile and diverse group belonging to the *Enterobacteriaceae* family and includes the ubiquitous *Klebsiella variicola* (Rosenblueth *et al*., 2004; Barrios-Camacho *et al*., 2019). The latter is commonly isolated as part of plant microbiomes as it can promote plant growth by nitrogen fixation (Pinto-Tomás *et al*., 2009). It is also an emerging human pathogen (Rodríguez-Medina *et al*., 2019) causing urinary tract infections (Potter *et al*., 2018), as well as in other diseases in wild and farm animals (Giannattasio-Ferraz *et al*., 2022). However, *K. variicola* has been largely neglected due to a historic taxonomic misclassification (Martínez-Romero *et al*., 2018; Rodrigues *et al*., 2018). Recent developments including fast multiplex PCR (Garza-Ramos *et al*., 2015), mass spectrometry profiles (Rodrigues *et al*., 2018), and phylogenomic analyses (Lam *et al*., 2021) allowed the appropriate *K. variicola* identification and prompted studies addressing this species’ biology, population structure and virulence determinants (Potter *et al*., 2018). Analyses of the *K. variicola* pangenome revealed the existence of nine different pili, including two well-studied chaperon usher systems type 1 fimbriae encoded in the *fim* operon and the type 3 fimbriae (T3F) encoded by *mrk* operon (Potter *et al*., 2018).

To study how morphotypic diversity emerges and how it is maintained in *Klebsiella pneumoniae species complex,* and more specifically in *K. variicola*, we used a previous evolution study (Nucci, Rocha and Rendueles, 2022), in which we evolved in parallel two hypervirulent *K. pneumoniae* strains (Kpn NTUH and Kpn BJ1) and one environmental *K. variicola* strain (Kva 342) as well as their non-capsulated isogenic mutants. The non-capsulated mutants were generated by an inframe deletion of *wcaJ*, the first gene of the biosynthetic pathway and the gene most commonly mutated in lab-evolved non-capsulated clones (Buffet, Rocha and Rendueles, 2021) and genomic datasets (Haudiquet *et al*., 2021). We propagated these six different genotypes for 675 generations in two nutrient-rich environments (artificial sputum and LB), and three nutrient-poor environments (artificial urine, soil and minimal media supplemented with glucose). Prior to each daily transfer, populations were re-homogenized by vigorous pipetting, and a bottleneck of 1% was applied. The described evolution experiment allowed us to show that originally capsulated populations rapidly diversified generating several morphotypes, all of which relied on the capacity of each clone to produce the polysaccharidic capsule. On the one hand, we observed small and translucent non-capsulated colonies, most often generated by IS insertions or point mutations in *wcaJ*, the initial glycosyltransferase and first enzyme of the capsule biosynthesis pathway. On the other hand, large, bulky and hypermucoviscous capsulated colonies also evolved frequently due to point mutations in *wzc*, another gene involved in capsule synthesis (Nucci, Rocha and Rendueles, 2022). However, morphological diversification of the originally non-capsulated populations was rarer and remained to be addressed.

Here, we followed throughout the evolution experiment the emergence of a novel rough and dry morphotype (i.e. rdar-like) in non-capsulated strains, similar to the rdar-like morphotypes of other Enterobacteria, including *E. coli* and *Salmonella* (Römling, 2005; Cimdins *et al*., 2017). We identified its genetic basis, quantified its fitness effects, and determined the underlying selection forces. Finally, we searched in the literature and in a large genomic dataset for the prevalence of the identified mutations leading to this morphotype in natural populations.

Our work revealed that novel morphotypes under negative frequency-dependent selection emerged in non-capsulated populations and mimic those found in the clinic.

## RESULTS

### Emergence of rdar-like morphotypes in non-capsulated populations of *K. variicola*

The regular plating of independent populations during the evolution experiment (see Materials and Methods) revealed the emergence of colonies displaying peculiar morphologies in six independent populations of non-capsulated *Klebsiella variicola* 342, *i.e.* twenty percent of all non-capsulated Kva 342 populations (Figure 1A). Such morphotype was not observed in populations descending from the capsulated ancestor nor from the two *Klebsiella pneumoniae* strains, Kpn NTUH or Kpn BJ1. These morphotypes evolved thrice in M02 and thrice in AUM, two environments with low-carrying capacity, but not in ASM or LB, two environments with high-carrying capacity. We isolated one clone of each population for further analyses. Despite the similarities of the morphotypes at the single-colony level (originating from single cells), at the population-level (originating from an overnight culture), a morphological difference was evident between 4D2, 4D6 and 6B3 (morphotype 1) vs 4D4, 6B1 and 6B4 (morphotype 2) clones (Figure 1A).

**Figure 1.**
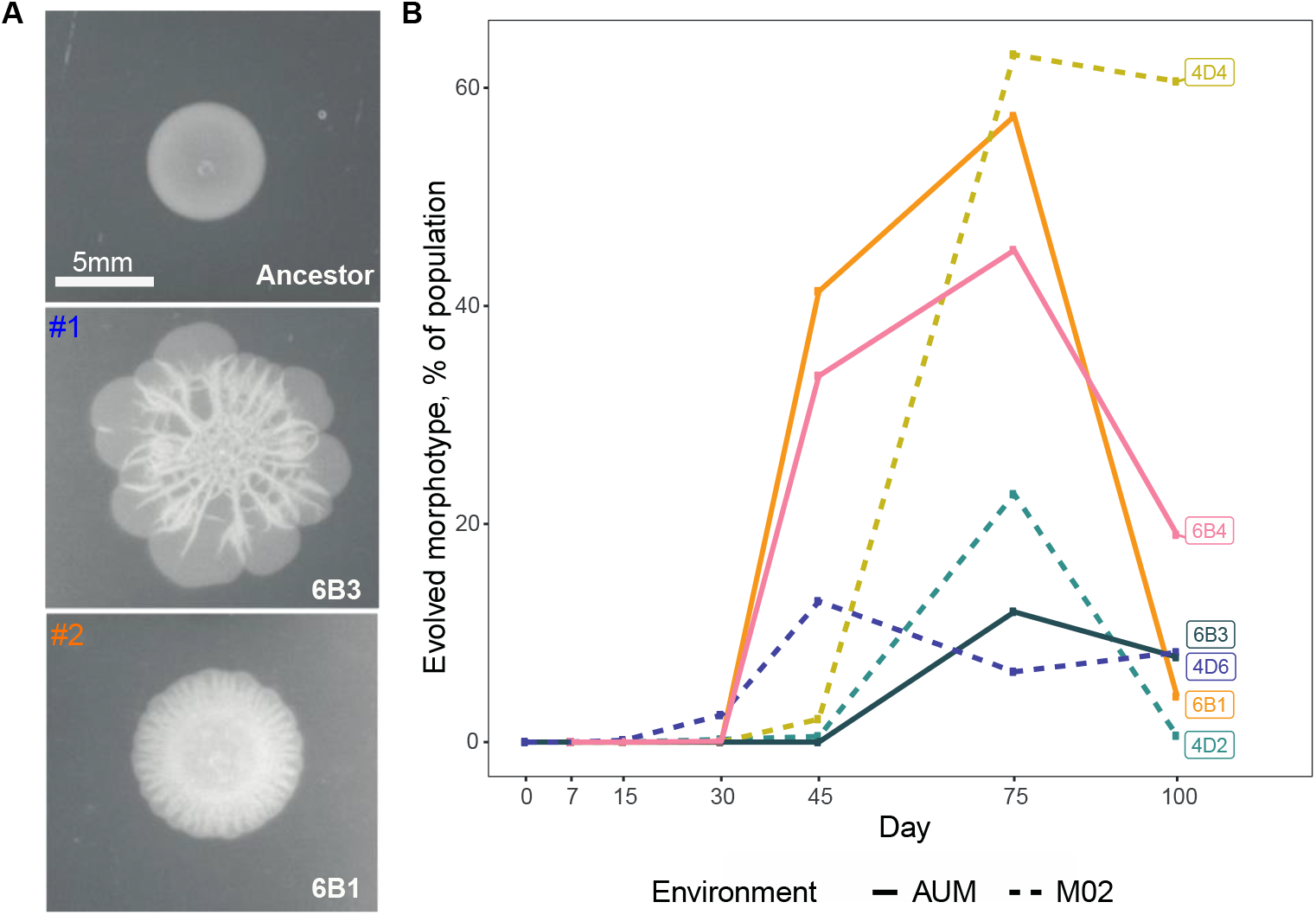
Phenotype and evolution of rdar- like morphotype in *K. variicola* populations. **A**. Representative images of the ancestor and two rdar-like population-level colonies from clone 6B3 (morphotype 1) and 6B1 (morphotype 2). Five µL of an overnight culture were spotted on an LB plate and allowed to grow for 24 hours. **B**. Population dynamics of evolved morphotypes in the six independent populations. Populations with morphotype 1 (4D2, 4D6 and 6B3) are displayed in cold colors (blue), whereas populations belonging to morphotype 2 (4D4, 6B1 and 6B4) are depicted with warmer colors (orange).

To determine the frequency of the morphotypes and the population dynamics, we plated the six evolving populations at all intermediate time-points. After 50 cycles, *ca* 330 generations, the morphotypes had already emerged in all populations. Clones displaying morphotype 2 reached significantly higher maximum frequencies compared to those of morphotype 1(t-test, *P* = 0.009). These ranged between 12.7 and 66.6% of the population (Figure 1B). However, after reaching maximum frequency, most morphotypes experienced a strong decrease in frequency, and towards the end of the experiment reached a frequency of ∼16% of the total population.

The observed morphotypes are similar to the rdar phenotype observed in other enterobacteria, including *E. coli* and *Salmonella* (Römling, 2005), or the abovementioned wrinkly spreader of *Pseudomonas fluorescens* (Rainey and Travisano, 1998). These well-known morphotypes rely on the expression of specific surface adhesins and exopolysaccharides, most notably curli and cellulose (White and Surette, 2006; Cimdins *et al*., 2017; Di Sante *et al*., 2018). A bioinformatic search for the *csgA-G* operon responsible for curli formation in the genome of Kva 342 revealed no match (see Methods, Table S1). We then looked for the *bcsABCDEFGQZ* operon responsible for cellulose biosynthesis. The Kva 342 had a complete cellulose synthesis. We tested the ability of the strain to produce cellulose by the specific binding to calcofluor dye. No differences in colony morphotype were observed between the ancestor and the evolved clones (data not shown). We thus conclude that the emergence of these two novel rdar-like morphotypes in *K. variicola* does not rely on the expression of curli or exopolysaccharides, the two known mechanisms involved in the formation of rdar-like morphotypes in Enterobacteria (Römling, 2005; White and Surette, 2006)

### Mutations in two type 3 fimbriae regulators are responsible for the rdar-morphotypes

To determine the genetic basis of the rdar-like morphotype, we performed whole-genome sequencing of four randomly-chosen rdar-like clones from distinct populations, two of each morphotype. We then compared it to the ancestral sequence of Kva 342 (Table S2). We observed a remarkable mutational convergence in two genes *mrkH* (morphotype 1) and *nac* (morphotype 2) (Table 1). Both code for nucleic acid binding proteins, recognizing either RNA (confidence score = 0.85) and DNA (confidence score = 0.98), respectively, as predicted by DeepFRI (Gligorijević *et al*., 2021). Indeed, MrkH binds to the α-CTD of RNA polymerase and acts as a transcriptional activator of the *mrk* operon involved in the synthesis of T3F (Wilksch *et al*., 2011). NAC is a LysR-type transcriptional activator known to be expressed in nitrogen-limited conditions and activating sigma70-dependent genes (Bender, 1991). PCR and sanger sequencing of the two non-sequenced clones (4D2 and 6B1) also revealed mutations in *mrkH* and *nac* respectively (Table 1). Interestingly, a search for the 15-bp consensus sequence for the LysR-binding box in *Klebsiella* (ATA-N9-TAT) (Frisch and Bender, 2010) revealed two hits in the mrk operon; one 200bp upstream the start codon of *mrkA* and a second one upstream the promotor of *mrkJ* (Figure S1). Interestingly, this LysR-binding box, overlaps the ‘mrkH box’, a palindromic sequence to which MrkH binds to control transcription of *mrkHI* and *mrkABCDF* clusters. Deletion of the ‘mrkH box’ also leads to reduction of *mrkJ* expression (Ares *et al*., 2017). This suggests that mutations in either *mrkH* or *nac* ultimately impact type 3 fimbriae.

**Table 1.** Convergent mutations in clones displaying rdar-like morphotypes.

| Clone | Morphotype | Environment | Position | Mutation | Change | Annotation |
| --- | --- | --- | --- | --- | --- | --- |
| 4D2 | 1 | M02 | 844 700 | IS insertion | 608/705 nt | *mrkH* |
| 4D4 | 2 | M02 | 5 497 313 | C→T | G275S | *nac* |
| 4D6 | 1 | M02 | 844 713 | IS insertion | 621/705 nt | *mrkH* |
| 6B1 | 2 | AUM | 5 498 042 | C→T | R228C | *nac* |
| 6B3 | 1 | AUM | 845 042 | G→A | P98S | *mrkH* |
| 6B4 | 2 | AUM | 5 498 183 | C→A | intergenic (-48/+212) | *nac /argH* |

To test whether the rdar-like morphotype was dependent on the *mrkH* and *nac,* we restored the ancestral allele in four different evolved clones from different environments. Restoration of the ancestral mrkH allele resulted in the loss of morphotype 1 (Figure S2A), whereas restauration of *nac* allele in clones 4D4 and 6B1 also resulted in the loss of morphotype 2 (Figure S2B). We then showed that SNPs in *mrkH* or *nac* were sufficient to produce the evolved morphotypes (Figure S2C) when inserted in an ancestral genotype. Finally, to show that IS insertions in the 3’ of the gene, due to IS903 from IS5 family in *mrkH* were equivalent to a loss of function, we generated an in-frame deletion of the entire *mrkH* gene. This led to the emergence of the rdar-morphotype (Figure S2C). Overall, our results show that mutations in known regulators of T3F are responsible for the rdar-like morphotype.

### Mutations in *mrkH* and *nac* decrease aggregation and limit biofilm formation but increase growth rate

Rdar-like morphotypes are well known for having altered intercellular interactions which result in differences in aggregation and formation of biofilm (Da Re and Ghigo, 2006). We tested this and showed that evolved clones had a diminished capacity to aggregate compared to the ancestor (Figure S3A). As expected, this was mostly associated with a decreased capacity to form biofilm except for the evolved clones 4D2 and 4D6, both of which had an IS inserted in *mrkH* (Figure S3B).

We hypothesize that mutations in *mrkH* and *nac* could be either responsible for the decreased aggregation, for changes in biofilm formation or for both. We tested this using the abovementioned mutants. To quantify aggregation, we measured the absorbance of the top layer of sitting cultures through time as a proxy for sedimentation. In aggregation tests, higher relative absorbance represents low sedimentation. Biofilm formation was measured using the traditional crystal violet staining method. The reversion of SNPs in *nac* to the ancestral allele restored wild type levels of both aggregation (Figure 2) and biofilm formation (Figure 2C). Insertion of one SNP resulting in the amino acid change G275C in the ancestral background recapitulated the evolved phenotype (Figure 2C). We thus conclude that SNPs in *nac* alone are enough to explain the changes observed in the evolved morphotype 2.

**Figure 2.**
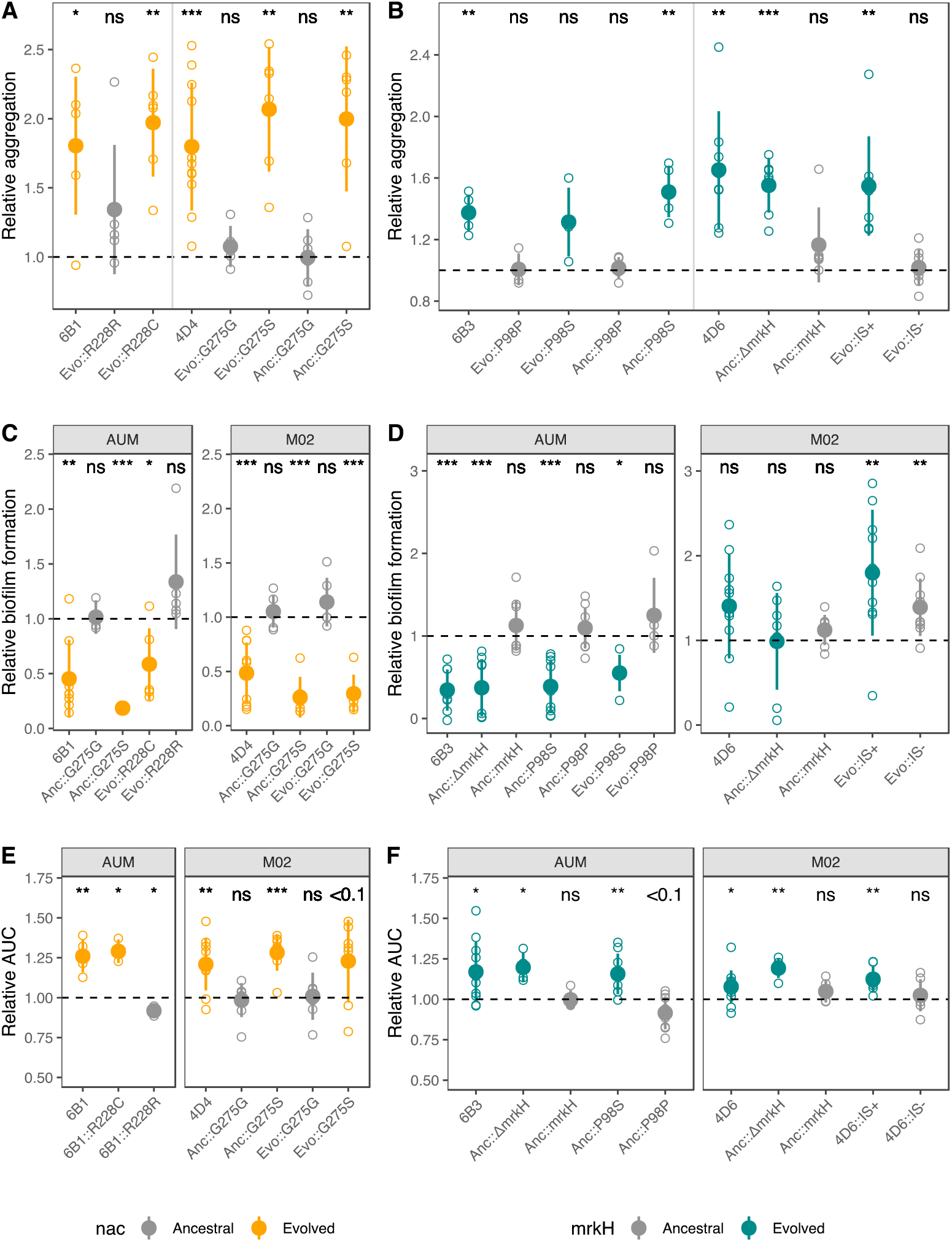
Mutations in *mrkH* and *nac* diminish aggregation biofilm formation and increase growth. **A.B.** Relative aggregation of *nac* (**A**) and *mrkH* (**B**) mutants is quantified by the absorbance (OD600) of the top layer of culture in static conditions during 24 hours and the results are the ratio of the values for evolved over the initial clones. Reported data correspond to differences after 4.5 hours. High absorbance results from low aggregation levels. Calculation of the area under the aggregation curve results in qualitatively similar results. **C. D.** Biofilm formation of *nac* (**B**) and *mrkH* (**D)** mutants was measured in the evolution growth media in which each mutation emerged. **E.F.** Area under the growth curve of evolved and ancestral *nac* I and *mrkH* (**F**) alleles. Data is represented relative to the ancestral strain (dashed line). The AUC was calculated using the formula *trapz* from the pracma package for R. Grey points represent clones with ancestral alleles whereas colors indicate clones with evolved *mrkH* (green) or *nac* (orange) alleles. Small open points indicate independent biological replicates. Large closed points represent the average of biological replicates and error bars indicate the standard deviation. Statistical analysis was performed to compare all alleles to its non-capsulated ancestor. One-sample two-sided t-test, difference from 1. * P<0.05, **P<0.01, *** P<0.001, ns P≥0.05.

Reversal of the amino acid change P98S in the gene *mrkH* or reversal of the IS insertion to the ancestral alleles restored wild type aggregation. Similarly, insertion of the evolved allele or deletion of *mrkH* in the ancestor recapitulated the evolved phenotype and decreased aggregation (Figure 2B). Absence of significant differences between the deletion of the gene *mrkH* and P98S in the ancestral background indicated that this amino acid change results in a loss of function (Two-sided t-test, P > 0.05). However, we do observe interesting differences in biofilm formation, depending on the environment. As expected, in AUM, *mrkH* mutations alone could explain the differences between evolved and ancestral phenotypes. This was not so in M02. For instance, the deletion of *mrkH,* or the reconstitution of a full *mrkH* in the evolved clone did not significantly alter biofilm formation compared to their wild type or the IS-interrupted genes, respectively. This suggests that *mrkH* is not being selected for changes in biofilm formation in our evolution experiment, but rather for changes in aggregation.

We then tested whether *nac* and *mrkH* mutations in evolved clones resulted in increased growth rate, as we would expect due to adaptation to the environment. We calculated the area under the growth curve (AUC), a measure that takes into account the lag time, maximum growth rate and population yield. All evolved clones (Figure S3C), as well as those with the insertion of *mrkH* or *nac* evolved allele (or deletion of the gene) in an ancestral background, had a growth advantage irrespective of the environment (Figure 2EF). Overall, the differences in growth observed in *mrkH* and *nac* mutants suggest that these mutations are adaptive.

Finally, to test if phenotypic effect in *mrkH* mutants results from changes in the expression of T3F, we performed quantitative RT-PCR. Loss-of-function of *mrkH* reduced significantly the expression of *mrkA* (the major subunit pilin of T3F) in both AUM and M02 (Two-way ANOVA, dF= 5, F = 4.8, *P* = 0.007) (Figure S4).

Taken together, mutations in *mrkH* and *nac* are responsible the decreased aggregation and faster growth of evolved rdar-like morphotypes. Further, mutations in *mrkH* also diminish biofilm formation, but in an environment-dependent way, suggesting that its roles in biofilm formation was not the primary force selecting for this morphotype.

### Mutations in *mrkH* are adaptive and under negative frequency-dependent selection in AUM

The parallel emergence of rdar-like morphotype in populations that evolved in M02 and AUM suggested that it is adaptive. To test this hypothesis, we focused on the clones with mutations in *mrkH*. We excluded *nac* in this analysis because it belongs to one of the largest family of regulator proteins known and could alter numerous other phenotypes via activation of sigma70-dependent genes (Bender, 1991). As expected, evolved clones had a significant fitness advantage compared to the ancestor in both environments (Figure S5A). We then performed direct competitions in a 1:1 ratio of the ancestral strain against its isogenic P98S or the loss-of-function *mrkH* deletion in the two environments. Additionally, we also competed an evolved clone against its isogenic mutant in which the ancestral *mrkH* was restored. In AUM, mutations in *mrkH* are enough to explain the increase in fitness of the evolved clones (Figure 3AB). In M02, *mrkH* increases fitness in an evolved background, but not in an ancestral background. This suggests that it requires other mutations to improve fitness (Figure 3AB).

Some rdar-like morphotypes reached high frequencies during the evolution experiment, but towards the end, their frequencies decreased. This could be indicative of negative frequency-dependent selection. To test this, we repeated the competitions experiments but with the evolved alleles in in an initial ratio of 1:9. In M02, fitness gains were equivalent across the different inoculation ratios (Two-sampled t-test, *P* = 0.71, N=3). However, in AUM, we found that when inoculated in the minority, evolved alleles were not only fitter than the ancestor (Figure 3CD, Figure S4B), but the fitness gain was significantly higher, compared to competitions in which both clones were inoculated at similar frequencies (Two-sampled t-test, *P* = 0.04, N=3). Further, fitness of evolved allele correlated negatively with proportion at the beginning of the coculture (Figure 3E), in AUM but not in M02. Finally, even when inoculated in the minority, rdar-like morphotypes were more frequent at the end of the competition.

**Figure 3.**
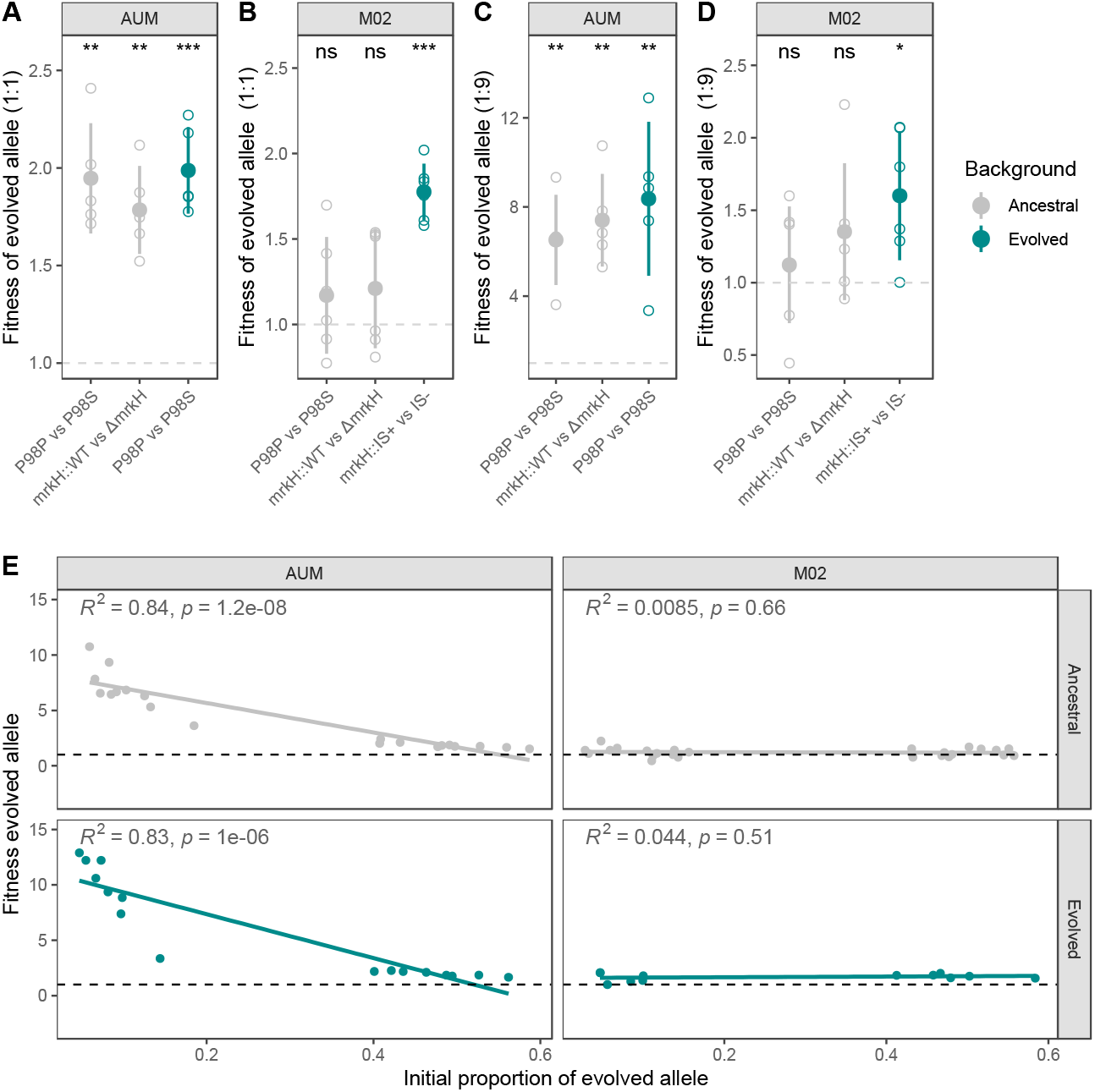
Fitness effects of mutations in *mrkH*. Competitions were performed at an initial ratio of 1:1 (**A, B**) and of 1:9 (**C, D**), with the ancestral allele in the majority. Grey dots correspond to competitions in which the ancestor was competed against isogenic mutants with carrying either a SNP in *mrkH* or a *mrkH* deletion. Green dots correspond to competitions among evolved clones which were reverted or not to ancestral allele. Each allele was tested in its respective evolutionary environment. Large closed points represent the average of biological replicates and error bars indicate the standard deviation. Open points indicate individual biological replicates One-sample two-sided t-test, difference from 1. * P<0.05, **P<0.01, *** P<0.001, ns P>0.05. **E.** Correlation between fitness of evolved clone versus the initial proportion of the population. Each dot represents an independent biological replicate of an evolved:ancestral competition, across different genetic backgrounds. In grey evolved alleles were introduced in an ancestral background, whereas green represent competition against a reversion of the evolved towards an ancestral allele. P-values were calculated and plotted using the stat_cor function in the *ggpubr* package.

Taken together, *mrkH* mutations result in reduced *mrkA* expression and reduced aggregation. In M02, they are adaptive in the genetic context in which they emerged, whereas, in AUM, these mutations are adaptive even in the ancestral background and are under negative-frequency-dependent selection.

### The presence of the capsule limits effect of *mrkH* mutations on aggregation

The emergence of the rdar-like morphotype was exclusively observed in the non-capsulated background. We first hypothesized this could be due to potentially deleterious effects for the cell in a capsulated background. However, *mrkH* mutant clones (both gene deletions and substitution with evolved allele) were obtained easily thereby suggesting that these clones are not deleterious and could have emerged in the experiments with capsulated strains. Of note, insertion of *mrkH* and *nac* mutations produced a very noticeable morphotype, which could have gone unnoticed (Figure S6A). The resulting colonies were slightly rough but not dry, and thus, strictly speaking they are not rdar-like, as the capsule masks the dryness, but roughness at the borders can be observed. Next, we hypothesized that the *mrk* operon was not expressed in our evolutionary conditions in capsulated background and was therefore “invisible” to selection. But qRT-PCR showed marginal or no differences in expression of the *mrkA* between capsulated and non-capsulated genetic background (Figure S6B). We then hypothesized that mutations in *mrkH* had milder fitness effects, if any, on capsulated cells and were thus not strongly selected for. Direct competitions between the capsulated ancestor and evolved *mrkH* alleles showed that the latter were fitter. This was significantly so in AUM, both at 1:1 and at 1:9 ratio (Figure 4A). Comparison of fitness effects across capsulated and non-capsulated clones showed that these effects were only marginally larger in non-capsulated than in capsulated clones. This suggests that rdar-like mutations could be under weaker selection in capsulated clones (Figure S6C).

**Figure 4.**
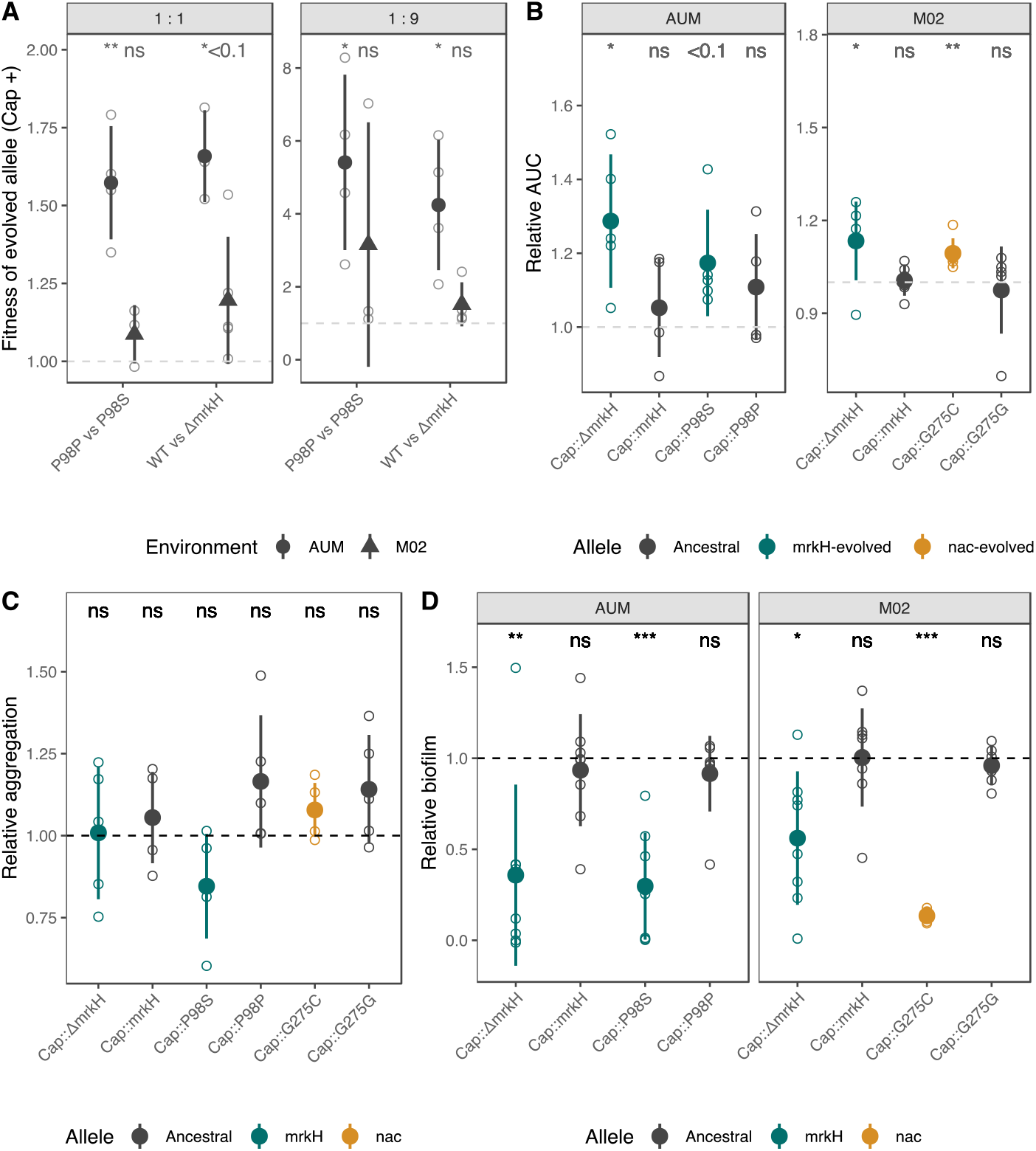
Effects of mutations in capsulated ancestors. **A.** Fitness effects of evolved alleles in capsulated strains when mixed in a 1/1 (0.5à and in a 1:9 (0.1) ration. **B.** Area under the growth curves relative to the capsulated ancestor. **C.** Aggregation of *mrkH* and *nac* mutants relative to the capsulated ancestor. Reported data correspond to differences after 4.5 hours in absorbance (OD600). **D.** Biofilm formation of *mrkH* and *nac* alleles was measured in the evolution growth media in which each mutation emerged and compared to the non-capsulated mutants (dashed line). Black points represent capsulated clones with ancestral alleles whereas dark green points indicate capsulated clones with evolved *mrkH* (green) or *nac* (dark orange) alleles. Small open points indicate independent biological replicates. Large and closed points represent the average of biological replicates and error bars indicate the standard deviation. Statistical analysis was performed to compare all alleles to its non-capsulated ancestor. One-sample two-sided t-test, difference from 1. * P<0.05, **P<0.01, *** P<0.001, ns P>0.05.

To test which phenotype is driving the selection for rdar-like clones in non-capsulated, we tested growth, aggregation and biofilm formation in a capsulated background (Figure 5BCD) and compared it to the respective non-capsulated construction (Figure S6DEF). Growth curves of capsulated mutants revealed that the relative effect of ancestral and evolved *mrkH* and *nac* alleles (Figure 4B) is similar in capsulated and non-capsulated strains (Figure S6D). Of note, in M02, overall increased growth rate of capsulated clones is due to the large fitness advantage provided by the capsule in Kva 342 in M02 (Figure S6D) (Buffet, Rocha and Rendueles, 2021). These results imply that the absence of rdar-like morphotypes in capsulated strains are not due to skimmer advantage during growth. This further suggests that selection would be acting on another phenotype other than growth. Mutations in *mrkH* and *nac* decreased biofilm formation in capsulated strains (Figure 4D), in a similar magnitude than in the non-capsulated strains (Figure S6F). This again suggests that selection would not be acting on biofilm formation. However, we observed that in a capsulated background, evolved *mrkH* or *nac* alleles did not significantly impact aggregation. This is opposite to what it is observed in a non-capsulated background (Figure 2B). (Figure 4C). In fact, there is a significant difference in aggregation between ancestral and evolved *mrkH* and *nac* alleles in function of the presence of a capsule (Figure S6E).

**Figure 5.**
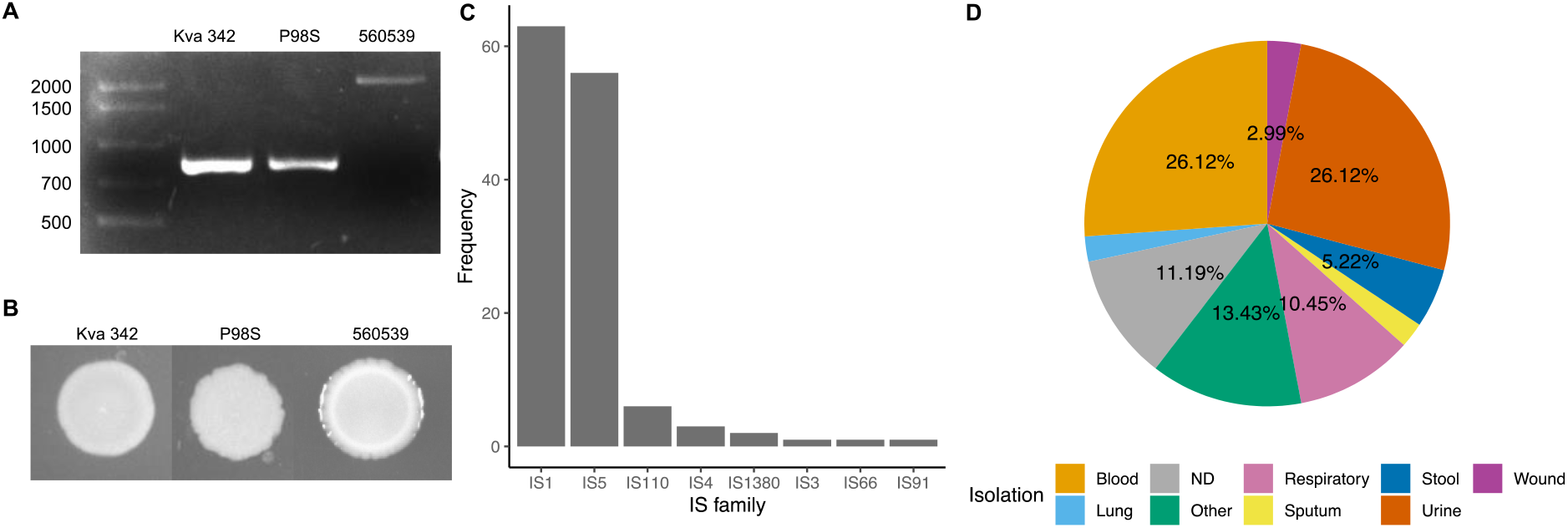
Interruption of *mrkH* gene by IS insertion in natural isolates. **A.** PCR confirmation that the *mrkH* truncation and the break in contig is due to the presence of an IS in the gene in strain MRSN-560539. **B.** Colony morphology of capsulated Kva 342, its isogenic *mrkH* mutant resulting in P98S and strain MRSN-560539. **C.** Family of ISs found to interrupt *mrkH* in genomes from the PATRIC database. **D.** The source of isolation of genomes in which an IS was identified co-localizing with *mrkH.* Metadata was retrieved from the PATRIC database and manually curated. Lung and sputum isolates represented less than 2% each.

Taken together, among the three traits we assessed (growth, aggregation and biofilm) that could directly contribute to fitness of the evolved rdar-like morphotype, only aggregation was different in capsulated clones when compared to non-capsulated. This strongly suggests that *mrkH* alleles are being selected for their impact in cell-to-cell aggregation.

### Truncations and IS insertions in MrkH are found in natural isolates

Recently, a similar rdar-like morphotype was described in a clinical isolate of *K. pneumoniae* from a blood sample (PBIO3459)(Sydow *et al*., 2022). As the rdar-morphotype evolved more readily in the non-capsulated background in our experiments, we first tested whether PBIO3459 could be non-capsulated. Using ISFinder (Siguier *et al*., 2006), we identified IS903 belonging to the IS5 family inserted in the *wza* gene (position 1127/1140), the outer membrane exporter of the capsule. Loss-of-function of Wza is associated to a loss of capsulation (Buffet, Rocha and Rendueles, 2021). We then searched the genome for MrkH and NAC protein sequences and compared them to those of our reference strains. The protein sequences of NAC are highly conserved between *K. pneumoniae* and *K. variicola*, revealing only one amino acid change between the two species in position 108. Accordingly, in PBIO3459, we only observed one additional amino acid change corresponding to N229K compared to the other *K. pneumoniae* proteins. Such change is one amino acid away from that of clone 6B1 (R228C) and is located in the DNA-binding domain (Figure S7). This could potentially explain the rdar-like morphotype. We then analysed the MrkH sequence. The MrkH sequence of PBIO3459 seemed truncated (Figure S7). We hypothesized that this could be the result of an IS insertion. Indeed, ISFinder revealed the presence of IS903 overlapping the end of *mrkH* sequence. Interestingly, IS903 is also responsible for mutations in *mrkH* in *K. variicola* 342. Independently of the mutation in *nac*, insertion of IS in *mrkH* could also explain the rdar phenotype in such clinical sample.

To further verify whether *nac* or *mrkH* mutations lead to a rdar-like morphotype in clinical isolates, we took advantage of a recently available *Klebsiella pneumoniae* panel of diverse clinical isolates (Martin *et al*., 2023). This collection consists of 100 sequenced *K. pneumoniae* clinical isolates worldwide and representative of the species diversity. We searched for NAC and MrkH protein sequences in the genomes. We detected one strain, MRSN-560539, isolated from urine, with a truncated *mrkH.* This truncated *mrkH* was found at the border of the contig. This could be by chance, but could also be suggestive of an IS insertion, as these often complicate genome assemblies. To test whether there was an anomaly in *mrkH* sequence, we performed a PCR and confirmed that an IS element, belonging to the IS21 family was inserted (position 428/729) (Figure 5A). We then analysed the colony morphology. We did not observe a dry colony, as the strain is capsulated (serotype KL36). However, we do observe roughness around the edges of the colony, similar to what was observed in the capsulated Kva 342 mutants we constructed (Figure 5B). NAC protein of this strain was identical to that of our wild type *K. pneumoniae* strains. This confirms that loss-of-function mutations in MrkH can also lead to a wrinkly phenotype in capsulated *K. pneumoniae* isolated in the clinic.

To test the broader prevalence of truncated *mrkH* gene or IS insertion events in MrkH, we downloaded and annotated the genomes (complete & draft) of all available *K. variicola* and 10 000 random *K. pneumoniae* genomes in the Pathosystems Resource Integration Center (PATRIC) genome database (Wattam *et al*., 2014). We searched for MrkH homologs and observed that ∼4% and ∼17% of MrkH proteins of *K. variicola* and *K. pneumoniae,* respectively, were either truncated or found at the border of contigs (Table S3) and could potentially exhibit rdar-like morphotypes. We then tested whether these proteins with abnormal lengths were interrupted by an IS. Indeed, we observed at least three *K. variicola* genomes had an IS inserted in MrkH (Table S4). In *K. pneumoniae,* we observed 133 (76 at the contig border and 57 inside the contig) of such events, most of which were mediated by IS5-like elements, among which IS903 (Figure 5C, Table S4). Isolates with interrupted MrkH were isolated from a human host, 35 of which (26%), were isolated from urine, as was strain MRSN-560539, and another 35 from blood cultures (Figure 5D).

Taken together our analyses suggest that the unique morphotype here described can be isolated in natural populations, including clinical isolates, and are mostly generated by IS sequences. Yet, these morphotypes are rare, as one would expect from traits that are under negative-frequency-dependent selection.

## DISCUSSION

We have been studying the outcome of a series of evolution experiments with three capsulated Klebsiella strains (and the non-capsulated mutant). This experiment has shed light on the general patterns of adaptation of the species independently of strain specificities, including the pervasive evolution of hypermucoviscosity (Nucci, Rocha and Rendueles, 2022). Similarly, it revealed how biofilm formation could evolve as a latent phenotype and how it coevolves with other fitness-related traits like surface-attached polysaccharides and population yield. This led to the identification of repeated mutations in the tip adhesin of T3F and revealed its conspicuous role in biofilm formation (Nucci, Rocha and Rendueles 2023).

Here, we focus on *K. variicola* and its morphotypic diversification. We expand the genomic analyses to a larger set of *K. variicola* and *K. pneumoniae* genomes to suggest that this morphotype is strongly associated to the urinary environment. Such morphotype is contingent on the genetic background and on the environment. Its emergence relies on the absence of bacterial capsule and on growth in nutrient-poor conditions. In addition, we identified that mutations in two regulators, *mrkH* and *nac*, were directly responsible for such morphotype. These mutations significantly reduce aggregation via decreased expression of surface fimbriae. Collectively, our data showed that the rdar-like morphotype in *K. variicola* is primarily selected due to its role in aggregation and not in growth or biofilm formation. This relies on several pieces of evidence. First, the increased growth rate of rdar-morphotypes which could easily explain increases in fitness and strong selective coefficient, is also observed in capsulated clones, where the morphotype is not selected for. Second, changes in the ability to form biofilm were environment-dependent, and thus could not explain the emergence of rdar morphotype in M02. The only trait which was consistent across environments and different across capsulated and non-capsulated populations was aggregation, suggesting this was the major selective force. Indeed, the fitness benefits of different *mrkH* mutations in capsulated clones are slightly less than in non-capsulated genetic backgrounds, especially in urine media (Figure S7C).

But why are non-aggregative phenotypes selected during the evolution experiment? Ancestral-like clones express T3F leading to clump formation and sedimentation. This can also lead to surface adhesion and biofilm formation at the bottom of the microtiterplate well. Rdar-like mutants have evolved to decrease expression of T3F, reduce costs associated to fimbriae production and grow faster (Figure 2). We foresee two main advantages for these ‘escape mutants’. First, our selection regime included vigorous pipetting previous to each transfer. Thus, larger, more cohesive cell clumps would be excluded in benefit of smaller, or isolated cells. Second, they will remain in suspension and access other resources, such as increased oxygen concentrations, which may be lacking at the bottom of the microtiterplate-well where other clones have fallen. Within a host, we speculate rdar morphotypes at low frequencies could favour dispersal of *Klebsiella* cells from a population during infectious episodes. It was recently shown that in the urinary tract, non-capsulated variants are often recovered (Ernst *et al*., 2020). This could constitute the first step for the evolution of rdar-like phenotypes prior to systemic dispersion by the blood, as observed in the clinical isolate PBIO3459 (Sydow *et al*., 2022).

Aggregation as a major driver of the rdar morphotype also helps understanding why during our evolution experiment such morphotypes emerged in non-capsulated populations of *K. variicola* but not in capsulated nor in *K. pneumoniae* populations. We hypothesize that for rdar-like morphotypes to provide a strong fitness advantage and be selected, some basal T3F-mediated aggregation in the population is required. In the case of capsulated clones, there is already very limited aggregation in the population due to masking of adhesins by the capsule (Schembri *et al*., 2005), thus loss-of-function mutations in *mrkH* does not strongly alter patterns of aggregation (Figure 5C). This also limits the fitness advantages compared to those in non-capsulated clones (Figure 5A and Figure S7). As for non-capsulated *K. pneumoniae*, we hypothesize that *mrkH* mutants did not emerge in our experiment due to subtle differences in T3F regulation across species. Deletion of *mrkH* significantly reduced but did not totally abolish *mrkA* expression in *K. variicola*, allowing for some T3F expression and aggregation. However*, mrkH* deletions in different *K. pneumoniae* strains resulted in undetectable level of *mrkA* mRNA and inability to form biofilm (Yang *et al*., 2013). Finally, the negative frequency-dependency also suggests that *mrkH* mutants provide larger fitness advantages only when rare, and thus require a majority of T3F-expressing cells.

In artificial urine, during competition experiments between rdar-like and wild type clones, we observed that even when inoculated in a minority, rdar-like clones not only increased their relative frequency, indicating higher fitness (Figure 4CD and S4), but they reached a majority (frequency > 0.5). This, together with the negative-frequency dependence advantage of the phenotype, could be suggestive of cheating. Cheating can be defined as a fitness relationship where a strain which performs poorly at a social trait in pure culture exploits another strain with high performance at the focal trait (Travisano and Velicer, 2004; Smith and Schuster, 2019). The effect of mixing would result in a relative within-group fitness advantage of the low-performance strain. This can be explained because the cheater cells do not pay their fair share of the cooperative act. Well-known cooperative acts are production of siderophores (West and Buckling, 2003), or fruiting body formation during multicellular development (Velicer and Vos, 2009). Here, if we consider that the main selective pressure in our experiment is aggregation and production of extracellular adhesins, *mrkH* mutants are unable to properly aggregate in isolation and could thus be *bona fide* cheaters. To the best of our knowledge, this is the first account of social exploitation in the *Klebsiella* genus.

Numerous evolution experiments performed in static environments or with some degree of spatial structure have revealed a remarkable morphotypic convergence in the emergence of rough, ‘wrinkly’ or rdar-like colonies. Formation of such colonies has been attributed to changes in exopolysaccharides (Spiers *et al*., 2002; Lin *et al*., 2022), or changes in expression of amyloid fibers like curli. Most of these morphotypes rely on regulatory pathways that respond to c-di-GMP levels (Blake *et al*., 2021). Indeed, this second messenger has been identified as an important determinant of diversification in *P. aeruginosa* biofilms (Flynn *et al*., 2016). Lastly, such morphotypes are very often frequency-dependent (Spiers *et al*., 2002; Poltak and Cooper, 2011; Udall *et al*., 2015). Indeed, they are stably present at low frequencies in the population, and their benefit is highest when they are rare. Such morphotypes also evolve in *K. variicola* and may also respond to c-di-GMP (via *mrkH,* (Wilksch *et al*., 2011)). Here, we show that they emerge by mutations in pathways independent of c-di-GMP (*nac*), and that their negative-frequency dependence is environment-specific. *K. variicola* morphotypes emerge independent of EPS or amyloid fiber production, but rather in changes in type 3 fimbriae. Most importantly, in most species these morphotypes are strongly aggregative allowing efficient colonisation of air-liquid interface (Poltak and Cooper, 2011; Udall *et al*., 2015; Blake *et al*., 2021) but in *K. variicola,* biofilm formation and aggregation is strongly impaired. Hence, even if these morphotypes are a common adaptive strategy across Bacteria, as shown by the incredible parallel evolution across microbial systems, they do not seem to be functionally convergent, as they seem to impact differently the ability to form biofilm.

This work further contributes to highlight the important role of insertion sequences in shaping genomes (Siguier, Gourbeyre and Chandler, 2014) and in the evolution of *Klebsiella*. For long, transposition of IS was associated to fitness loss. Here, we provide more evidence that IS can increase fitness and play an important role in adaptation (Consuegra *et al*., 2021; Frazão *et al*., 2022). More particularly, we pinpoint families IS1 and IS5, both carrying DDE transposases, as primary drivers of *Klebsiella* and its morphological diversification. IS903B from IS5 family is known to insert in *mgrB* and cause resistance to colistin, a last-resort antimicrobial peptide (Fordham, Mantzouratou and Sheridan, 2022). In our previous studies, we already showed that insertion of IS903B in the capsule operon was the main mechanism leading to the emergence of non-capsulated cells in capsulated populations of Kva 342 (Nucci, Rocha and Rendueles, 2022). Here, we observe that this can also extend to other morphological diversification mediated surface structures like fimbriae. Similar trends are also observed in *Klebsiella* natural populations. Large genomic analyses showed that at least 0.4% (3 out of 676) *K. variicola* and 1.4% (133 out of 9532) *K. pneumoniae* MrkH proteins were interrupted by an IS. This is most likely an underestimation as genes encoding MrkH were occasionally located at the border of the contig and a direct association to an IS insertion could not be determined. Yet, contig breaks are oftentimes an indication of an IS insertion, as we confirmed in strain MRSN-56039.

Taken together, our work provides important insight into the biology and evolution of *K. variicola,* a plant endosymbiont but also an emerging pathogen in cattle and humans, which has been significantly less studied than *K. pneumoniae sensu stricto*. Along our previous findings that one mutation in Kva 342 results in the *de novo* emergence of hypermucoviscosity (a proxy of hypervirulence) even in the absence of an immune system (Nucci, Rocha and Rendueles, 2022), this work further highlights the complexity of cell-to-cell and cell-to-surface interactions and how these evolve. Given the renewed interest for microbial products as fertilizers in agricultural settings to enable bacterial nitrogen fixation (Wen *et al*., 2021), and particularly of *K. variicola*, our study also contributes to critically advance our understanding of how this species evolves and responds to different environments, a requirement prior to any commercial exploitation.

## MATERIALS & METHODS

### Bacterial strains and growth conditions

i. *Strain*. *K. variicola* 342 (Kva 342, serotype K30) is an environmental strain, isolated from maize in the USA (Fouts *et al*., 2008). The K. pneumoniae diversity panel was acquired at BEI resources (https://www.beiresources.org/) and is available for research purposes under catalogue #NR-55604. *ii. Growth media.* AUM (artificial urine medium) was prepared as described previously (Brooks and Keevil, 1997). AUM is mainly composed of 1% urea and 0.1% peptone with trace amounts of lactic acid, uric acid, creatinine and peptone. ASM is composed of 0.5% mucin, 0.4% DNA, 0.5% egg yolk and 0.2% amino acids. LB is composed of 1% tryptone, 1% NaCl and 0.5% yeast extract. M02 corresponds to minimal M63B1 supplemented with 0.2% of glucose as a sole carbon source. *iii. Evolution experiment.* The evolution experiment was previously described (Nucci, Rocha and Rendueles, 2022). Briefly, six ancestral genotypes (Kva 342, Kpn NTUH and Kpn BJ1, and their respective non-capsulated mutants) were evolved in parallel for 100 days, accounting for *ca* 675 generations. To initiate the experiment, a single colony of each ancestral genotype was inoculated in 5 mL of LB and allowed to grow under shaking conditions at 37°C overnight. Twenty microliters of the diluted (1:100) overnight culture were used to inoculate each of the six independent replicates in the five environments. Each population was grown in a final volume of 2 mL in independent wells of 24 well microtiter plates. Every 24 hours, 20 μL of each culture was propagated into 1980 μL of fresh media and grown for 37°C under static conditions. Although each growth media had different carrying capacities, *i.e.* the maximum population size an environment can sustain, all cultures reached bacterial saturation in late stationary phase, ensuring that the different populations underwent a similar number of generations across media. Independently evolving populations were plated 28 times, every second day during ten days and every four days until the end of the experiment. Cross contaminations checks were routinely performed (Nucci, Rocha and Rendueles, 2022). *iv. Growth conditions.* Unless stated otherwise, experiments performed in the evolutionary environment were initiated by an overday culture in LB, an overnight culture in evolutionary media under shaking conditions. Pre-grown cultures were then diluted 1:100 into 1980 μL 24-well plates and allowed to grow for 24 hours at 37° under static conditions. *v. Primers.* Primers used in this study are listed in Table S5.

### Frequency of rdar-like morphotypes *mrk* mutants

To test the e frequencies of rdar-like morphotypes in the evolved populations, we aliquoted the glycerol stocks from days 7, 15, 30, 45, 75 and 100. Serial dilution was performed and plated on LB agar plates for CFU counting. The proportion of rdar-like clones was quantified.

### Whole genome sequencing and variant analyses

A single rdar clone from each population was isolated for whole genome sequencing. DNA was extracted from pelleted cells grown overnight in LB supplemented with 0.7mM EDTA with the guanidium thiocyanate method. Extra RNAse A treatment (37°C, 30min) was performed before DNA precipitation. Each clone was sequenced by Illumina with 150pb paired-end reads. Each evolved clone was compared to ancestral sequence using *breseq* (0.30.1) (Deatherage and Barrick, 2014) with default parameters.

### Mutant construction

Isogenic mutants were constructed by allelic exchange. i) *mrkH deletion.* 500bp upstream and downstream of the gene of interest were amplified. Cloning vector pKNG101 plasmid was also amplified using Phusion Master Mix (Thermo Scientific). Afterwards, pKNG101 was digested by DpnI (NEB BioEngland) restriction enzyme for 30 minutes at 37°C. Inserts and vector were then assembled using the GeneArt™ Gibson Assembly HiFi kit (Invitrogen), electroporated into competent *E. coli* DH5α strain and selected on Streptomycin LB plates (100 μg/mL for *E. coli*). Correct assemblies were checked by PCR. pKNG101 containing insert of interest was extracted using the QIAprep Spin Miniprep Kit then electroporated again into *E. coli* MFD λ-pir strain, used as a donor strain for conjugation in strains of interest. Single cross-over mutants (transconjugants) were selected on Streptomycin plates (200 μg/mL for *Klebsiella)* and double cross-over mutants were selected on LB without salt, supplemented with 5% sucrose, at room temperature. From each double-recombination, a mutant and a wild-type were isolated. Mutants were verified for their sensitivity to Streptomycin and by Sanger sequencing. ii) *Insertion and reversion* of evolved alleles was done in ancestor and evolved clone, respectively. The gene of interest (ancestor or evolved allele) was amplified using Phusion Master Mix (Thermo Scientific) and cloned into pKNG101 vector as described above. All mutants generated and used in this study are listed in Table S6.

### Biofilm formation

The capacity of a population or isolated clones to form a biofilm was performed as previously described (Buffet, Rocha and Rendueles, 2021). Briefly, each population or clones was pre-conditioned by allowing growth in LB overday, prior to inoculating overnight cultures in the environments in which the populations or clones evolved. Then, 20 μL of each overnight culture was inoculated into 1980 μL in 24-well microtiter plates and allowed to grow for 24 hours without shaking at 37°C. Unbound cells were removed by washing once in distilled water. To stain biofilms, 2100 μL of 1% crystal violet was added to each well for 20 minutes. The crystal violet was decanted and washed thrice with distilled water. The plates were allowed to dry under a laminar flow hood. Then, the biofilm was solubilized for 10 min in 2300 μL of mix with 80% ethanol and 20% acetone. Two hundred μL of each mix was transferred in a well of a 96-well plate. The absorbance of the sample was read at OD590.

### Aggregation test

An isolated colony was allowed to grow in 5 mL overnight in M02 medium at 37° under shaking conditions. Prior to the experiment, the absorbance (OD_600nm_) was measured and adjusted to OD_600_ = 2, and the cultures were transferred to static test tubes. Two hundred μL samples were sampled and the absorbance (OD_600nm_) was measured at defined time points (0; 1,5; 3; 4,5 and 24 hours) using an automatic plate reader Spark Control Magellan (TECAN). Samples were removed from the uppermost layer of tube cultures, roughly at the 4 mL mark. Decreasing absorbance represents the settling of agglutinated cell clumps. Calculation of the area under the aggregation curve with the function *trapz* from the R package pracma results in qualitatively similar interpretations. Values above 1 represent decreased aggregation compared to the ancestor.

### Fitness of *mrkH* mutants

To estimate the fitness advantage of mutations in *mrkH*, we performed direct competitions between the deletion, insertion and reversion mutants and their respective associated wild-types. Additionally, competitions between 6B3 and 4D6 evolved clones and the ancestor were performed. To initiate the competition experiments, individual clones were grown overnight in LB, and mixed in a 1 : 1 or 1 : 9 proportions. An aliquot was taken to estimate the initial ratio of each genotype by serial dilution and CFU counting as control of T_0_. The co-culture was then diluted 1 : 100 in 2 mL in the evolutionary environment in which the mutations emerged (AUM & M02). After 24 h of competition (T_24_) in 24-well microplate plates under static conditions, each culture was re-homogenized by vigorous pipetting and then serially diluted and plated. Rdar-like and wild-type colonies are clearly differentiated visually and counted separately. The competitive index of each genotype was calculated using the ratio 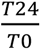. Competitions were only taken into account if initial frequencies for rdar-morphotype were in between 0.4-0.6 for 1:1 competitions and in between 0.02-0.18 for 1:9.

### Search for curli and cellulose operons

The proteic sequences of the experimentally validated curli biogenesis apparatus and the cellulose operon, both from *E. coli* were downloaded (accession numbers: Table S1). BlastP (v2.7.1+) with default parameters was used to search for each protein in the Kva 342 proteome. i). *Curli.* No hits were obtained (E-value < 10^-5^ & identity > 60%). ii). *Cellulose*. Each protein searched matched (E-value < 10^-5^ & identity > 60%) a protein in Kva 342 genomes. Further, these proteins were found in consecutive positions in the genome (*bcsGFEQABZC*).

### Search for MrkH proteins

The sequences for the *K. pneumoniae* diversity panel were downloaded on June 12 2023 from NCBI under BioProject PRJNA717739. All genomes corresponding to *K. variicola* in the Pathosystems Resource Integration Center (PATRIC) genome database (Wattam *et al*., 2014), filtered by good quality and text mined for *K. variicola* species (751 out of the 767), were downloaded on March 11 2022. Same procedure was applied for the first 10 000 genomes of *K. pneumoniae* (of which 239 were discarded). The genomes were checked for quality control and annotated with the pipeline PaNaCoTa (Perrin and Rocha, 2021) and the --prodigal option. Protein-Protein Blast (BLAST 2.7.1+) against either the Kva 342 MrkH was performed with the following option -max_target_seqs 100000 and an E-value smaller than 10^-5^. Sequences with an identity percentage of less than 90% were discarded. Reducing the threshold to 80% did not alter the number of sequences discarded.

### Quantitative RT-PCR

Twenty-four hour cultures in 24-well plates in the respective evolution environment were grown as abovementioned. RNA was extracted with the miRNA Extraction Kit (Macherey Nagel). Total RNA was measured and 400ng were used as matrix for cDNA amplification using the iScript cDNA Synthesis Kit. Samples were then treated for 30 minutes with DNase. Quantitive RT-PCR was performed using 1.5μL of amplified cDNA in a total volume of 15μL. As suggested by Gomes et al (Gomes *et al*., 2018), we used *rho* and *recA* as housekeeping genes. Data presented is based on calculations taking into account the geometric average of the two housekeeping genes. Taking one housekeeping gene or the other individually does not qualitatively alter any of the results.

## ACKNOWLEDGEMENTS

The authors would like to thank Katharina Schaufler and Stefan Heiden for providing the genomic sequence of strain PBIO3459. We also thank Samay Pande for helpful discussions during the writing of the manuscript. The sequencing of clones 4D4 and 6B4 was made at the Biomics Platform, C2RT, Institut Pasteur, Paris, France, supported by France Génomique (ANR-10-INBS-09) and IBISA.

## DATA AVAILABILITY

Raw reads are available at the European Nucleotide Archive (ENA), project number PRJEB54810. The sample names for rdar-like clones 4D4, 4D6 and 6B3 and 6B4 are ERS15562389, ERS12546754, ERS12546780 and ERS15562390 respectively.

All raw data generated in this study has been deposited in the public repository Figshare https://doi.org/10.6084/m9.figshare.23268791.

## AUTHOR CONTRIBUTIONS

OR conceived and designed the details of the study. AN, JJ and OR performed the experiments. AN and OR performed statistical analysis. OR performed the bioinformatics work, analyzed the data and wrote the manuscript. OR and EPCR secured funding, provided the resources and materials necessary for this study and revised the manuscript. All authors approved the final version of the manuscript.

## LEGENDS for SUPPLEMENTARY MATERIAL

**Figure S1. Genomic organization of the *mrk* operon in *K. variicola* 342.** Binding sequences were identified using the EMBOSS package *fuzznuc*, with default parameters (-auto) based on the published sequences of the MrkH box (Ares *et al*., 2017) and consensus LysR-binding site (Frisch and Bender, 2010).

**Figure S2. Mutations in *mrkH* and *nac* are enough to explain rdar-like morphotypes.** Colonies were grown overnight in LB. A five microliters droplet was deposited on LB agar and allow to grow at 37° for 24h. **A.** Colony pictures of evolved morphotype 1, as exemplified by the 4D6 and 6B3, as well as the restoration of the ancestral *mrkH* allele. A control colony resulting from the same double recombination event as the restored allele, but with an evolved allele, is also shown. **B.** Evolved morphotype 2 (4D4 and 6B1), as well as the restoration of *nac* allele, and its respective control colony. **C**. Insertion of the original non-capsulated ancestor of evolved SNPs in *mrkH* and *nac*, as well as clean isogenic deletions of *mrkH* gene.

**Figure S3. Aggregation (A), biofilm formation (B) and growth (C) of evolved rdar-like clones.** Data is represented relative to the non–capsulated ancestor (dashed line). Growth is expressed as AUC (area under the growth curve) and calculated using the formula *trapz* from the pracma package for R. Colors differentiate clones with mutations in either *mrkH* (green) or *nac* (orange). Small open points indicate independent biological replicates. Large closed points represent the average of biological replicates and error bars indicate the standard deviation. Statistical analysis was performed to compare all alleles to its non-capsulated ancestor. One-sample t-test, difference from 1. * P<0.05, **P<0.01, *** P<0.001, ns P>0.05.

**Figure S4. Expression of *mrkA* relative to the ancestor.** Expression of *mrkA* is expressed as the log_2_ of the fold change, compared to the non–capsulated ancestor. Small open points indicate independent biological replicates, each averaging at least two independent qPCR runs. Large closed points represent the average of biological replicates and error bars indicate the standard deviation. Statistical analysis was performed to compare all alleles to its non-capsulated ancestor. One-sample t-test, difference from 1. * P<0.05, **P<0.01, *** P<0.001, ns P>0.05.

**Figure S5. Fitness of evolved clones.** Competitions were performed with an initial ratio of 1:1 (**A**) and of 1:9 (**B**), with the evolved clones in the minority. Each clone was tested in its respective evolutionary environment. One-sample t-test, difference from 1. * P<0.05, **P<0.01

**Figure S6. Comparison between capsulated and non-capsulated mutants. A.** Colony morphotypes of capsulated Kva 342 with evolved *mrkH* (P98S) or *nac* (G275C) allele and a clean deletion of *mrkH* gene. **B.** Log_2_ expression difference of *mrkA* of capsulated genes using non-capsulated mutant strains as reference. **C**. Insertion of evolved SNPs in *mrkH* as well as clean isogenic deletions of *mrkH* gene in a capsulated ancestor in fitness at different inoculation ratios. **D.** Difference in the area under the growth curve between capsulated and non-capsulated mutants. **E.** Difference in aggregation between capsulated and non-capsulated mutants. **F.** Difference in biofilm formation between capsulated and non-capsulated mutants. Data are presented as the difference between capsulated and non-capsulated clones. One-sample t-test, difference from 0 (dashed line). * P<0.05, **P<0.01, *** P<0.001, ns P>0.05.

**Figure S7. Alignment of *K. variicola* and *K. pneumoniae* MrkH (A) and NAC (B) amino acid sequence.** MrkH and NAC sequences of Kva 342 and PBIO3459 (Sydow *et al*., 2022) were compared. Two other *K. pneumoniae* (Kpn BJ1 and NTUH) as well as another *K. variicola* CIP 80.47 were included as controls. Graphics and alignments were produced by http://multalin.toulouse.inra.fr/multalin/. Yellow arrows indicate SNPs in our evolution experiment and green breaks represent insertion of an IS in evolved clones.

**Table S1. Accession numbers of protein used as reference to search for the cellulose and curli biosynthesis operon in Kva 342.**

**Table S2. Mutations identified in all sequenced clones.** Differences between evolved and ancestral genomes were identified using *breseq* v 0.30.0 (Deatherage and Barrick, 2014) with default options.

**Table S3: Frequency table of protein length of MrkH.** MrkH homologs were identified with blastP. Results with an identity of more than 80% were kept and an e-value of < 10e-^5^. Wildtype MrkH is 234 amino acids.

**Table S4. Details of IS elements found in *mrkH* genes.** IS elements were identified using ISFinder (Siguier *et al*., 2006). Only hits with an e-value of <10e-5 were considered. For each IS found overlapping coordinates (including +100 or −100 bp) with those of MrkH, only the best hit per genome was taken into account.

**Table S5. Primers used in this study.**

**Table S6. Mutants used in this study.**

