## Supplementary Material for "Emergence of novel non-aggregative variants under negative frequency-dependent selection in *Klebsiella variicola*"

ORCID: AN, 0000-0001-7340-9075; EPCR, 0000-0001-7704-822X; OR, 0000-0002-6648-1594

### SUPPLEMENTARY FIGURES

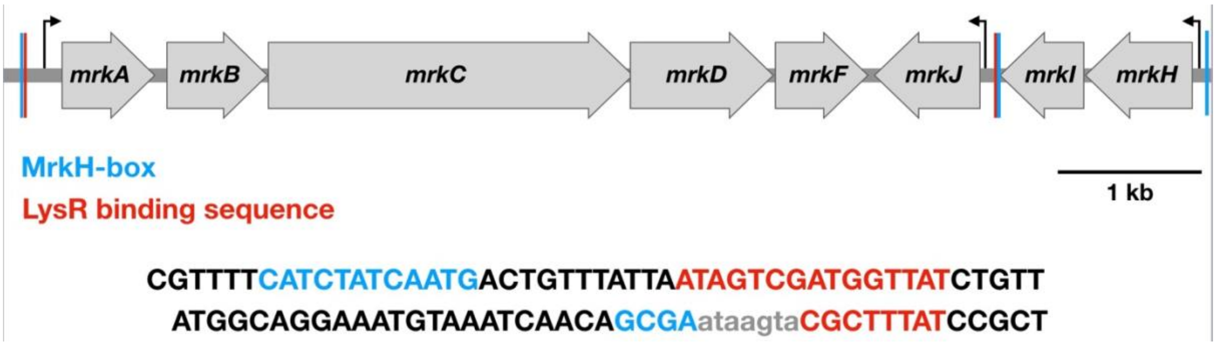


**Figure S1. Genomic organization of the *mrk* operon in *K. variicola* 342.** Binding sequences were identified using the EMBOSS package *fuzznuc*, with default parameters (-auto) based on the published sequences of the MrkH box (53) and consensus LysR-binding site (52).

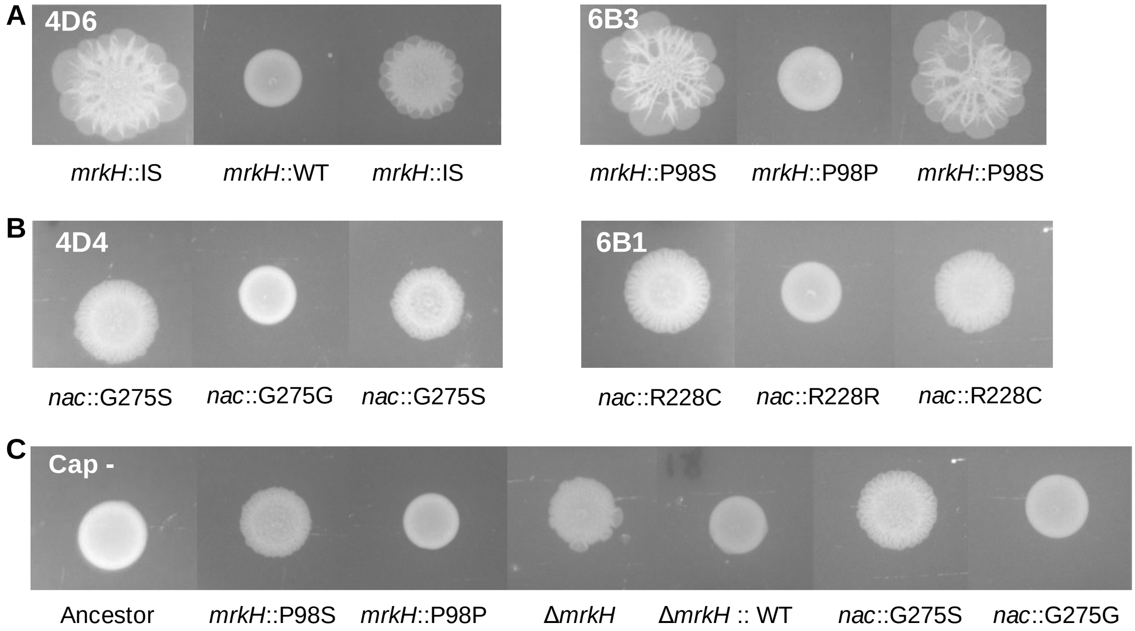


**Figure S2. Colony morphologies of mutants generated in this study**. Colonies were grown overnight in LB. A five microliter droplet was deposited on LB agar and allow to grow at 37° for 24h. **A.** Colony pictures of evolved morphotype 1, as exemplified by the 4D6 and 6B3, as well as the restoration of *mrkH* allele, and an evolved allele from the same double recombination event. **B.** Evolved morphotype 2 (4D4 and 6B1), as well as the restoration of *nac* allele, and an evolved allele from the same double recombination event. **C**. Insertion of evolved SNPs in *mrkH* and *nac*, as well as clean isogenic deletions of *mrkH* gene in the original non-capsulated ancestor.

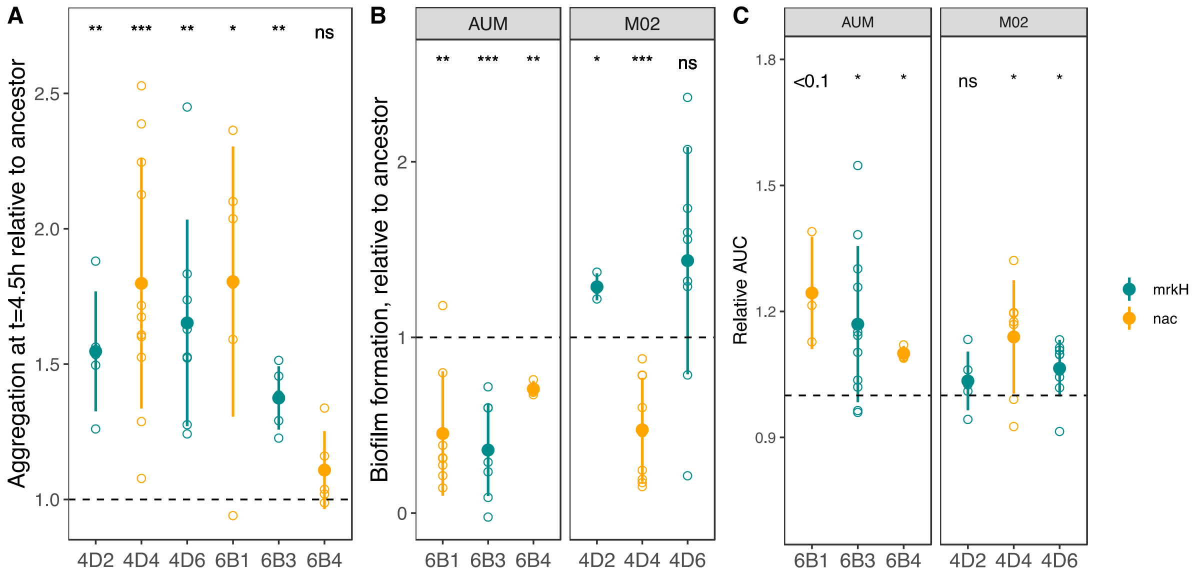


**Figure S3. Aggregation (A), biofilm formation (B) and growth (C) of evolved clones.** Data is represented relative to the non–capsulated ancestor **(**Dashed line). Growth is expressed as AUC (area under the growth curve), and calculated using the formula *trapz* from the pracma package for R. Colors differentiate clones with mutations in either *mrkH* (green) or *nac* (orange). Small open points indicate independent biological replicates. Large closed points represent the average of biological replicates and error bars indicate the standard deviation. Statistical analysis was performed to compare all alleles to its non-capsulated ancestor. One-sample t-test, difference from 1. * P<0.05, **P<0.01, *** P<0.001, ns P>0.05.

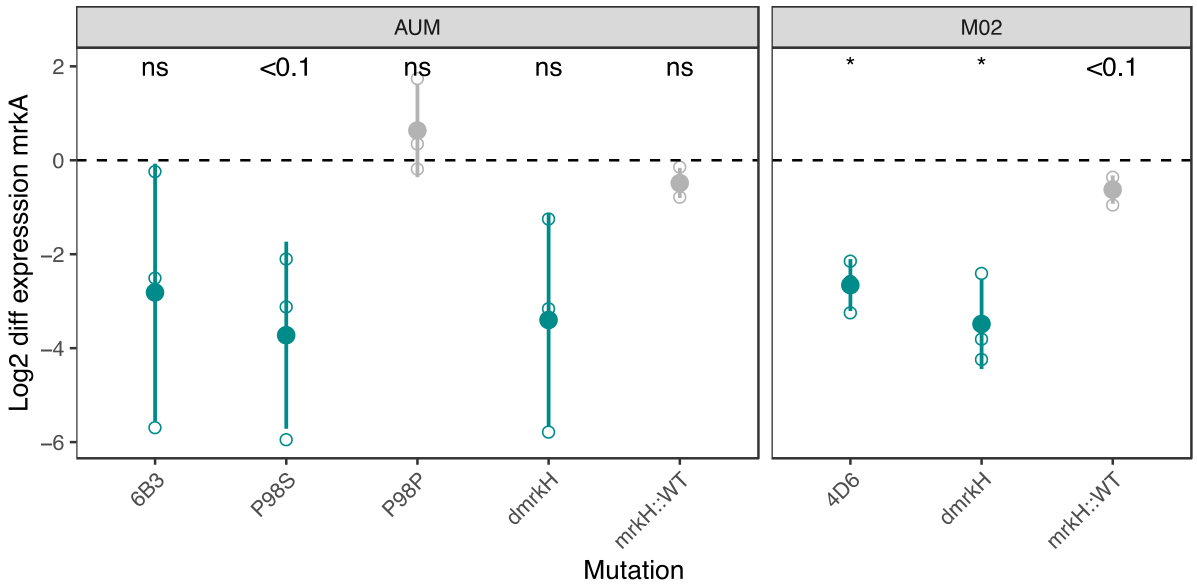


**Figure S4. Expression of *mrkA* relative to the ancestor.** Expression of *mrkA* is expressed as the $\log_2$ of the fold change, compared to the non–capsulated ancestor. Small open points indicate independent biological replicates, each averaging at least two independent qPCR runs. Large closed points represent the average of biological replicates and error bars indicate the standard deviation. Statistical analysis was performed to compare all alleles to its non-capsulated ancestor. One-sample t-test, difference from 1. * $P<0.05$, **$P<0.01$, *** $P<0.001$, ns $P>0.05$.

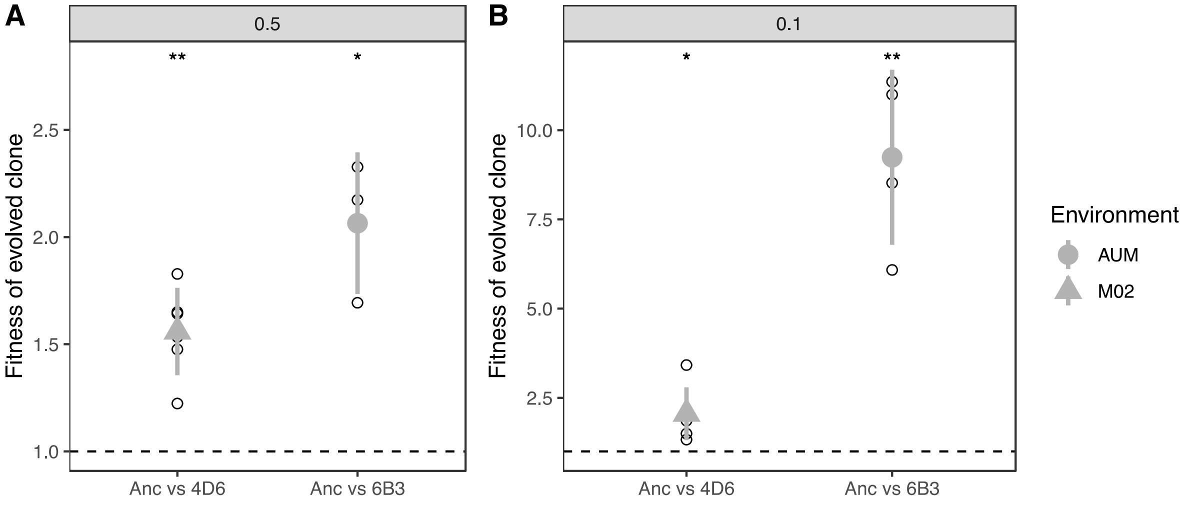


**Figure S5. Fitness of evolved clones.** Competitions were performed with an initial ratio of 1:1 (**A**) and of 1:9 (**B**), with the evolved clones in the minority. Each clone was tested in its respective evolutionary environment.

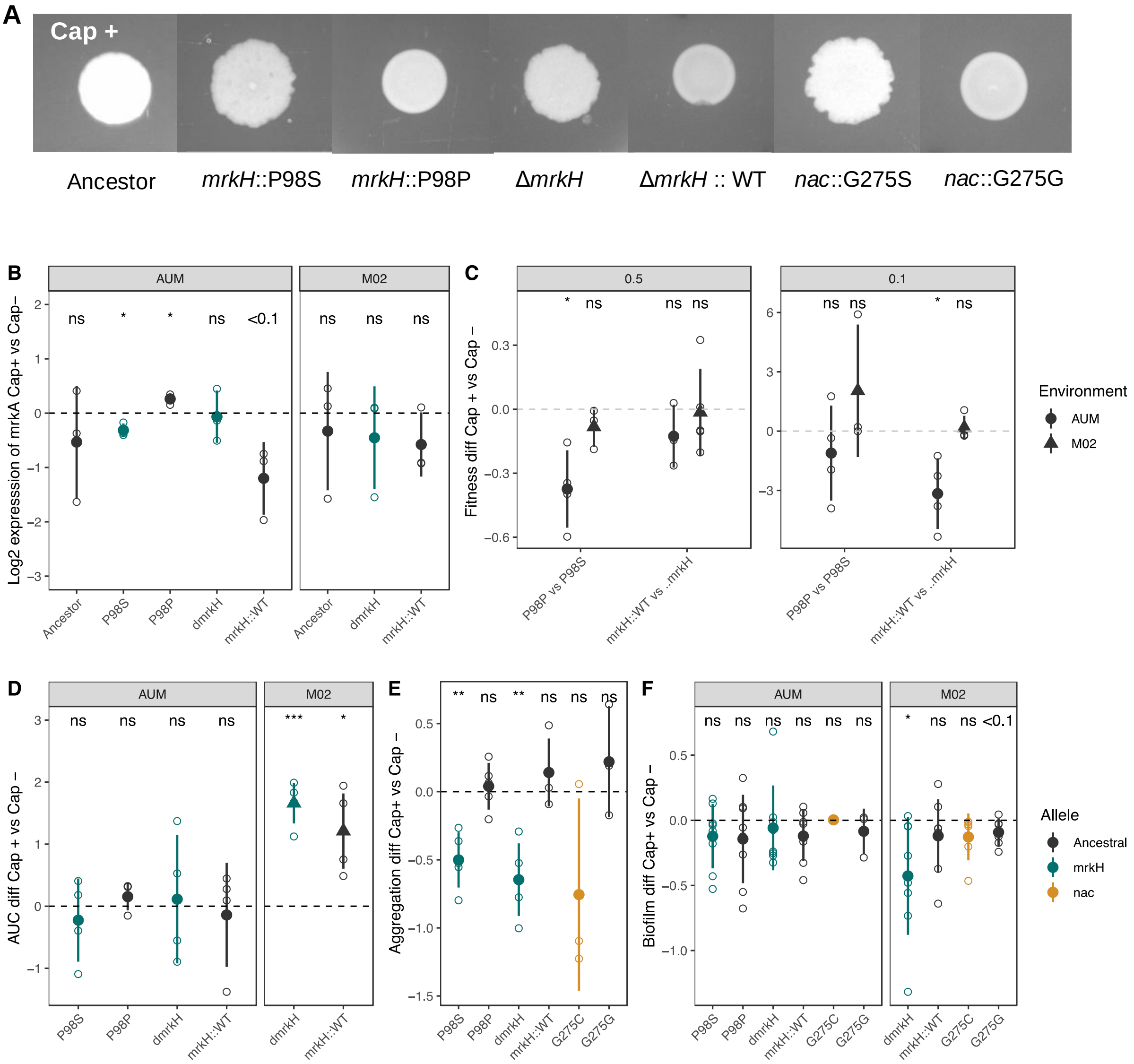


**Figure S6. Comparison between capsulated and non-capsulated mutants. A.** Colony morphotypes of capsulated Kva 342 with evolved *mrkH* (P98S) or *nac* (G275C) allele and a clean deletion of *mrkH* gene. **B.** $Log_2$ expression difference of *mrkA* of capsulated genes using non-capsulated mutant strains as reference. **C**. Insertion of evolved SNPs in *mrkH* as well as clean isogenic deletions of *mrkH* gene in a capsulated ancestor in fitness at different inoculation ratios. **D.** Difference in the area under the growth curve between capsulated and non-capsulated mutants. **E.** Difference in aggregation between capsulated and non-capsulated mutants. **F.** Difference in biofilm formation between capsulated and non-capsulated mutants.

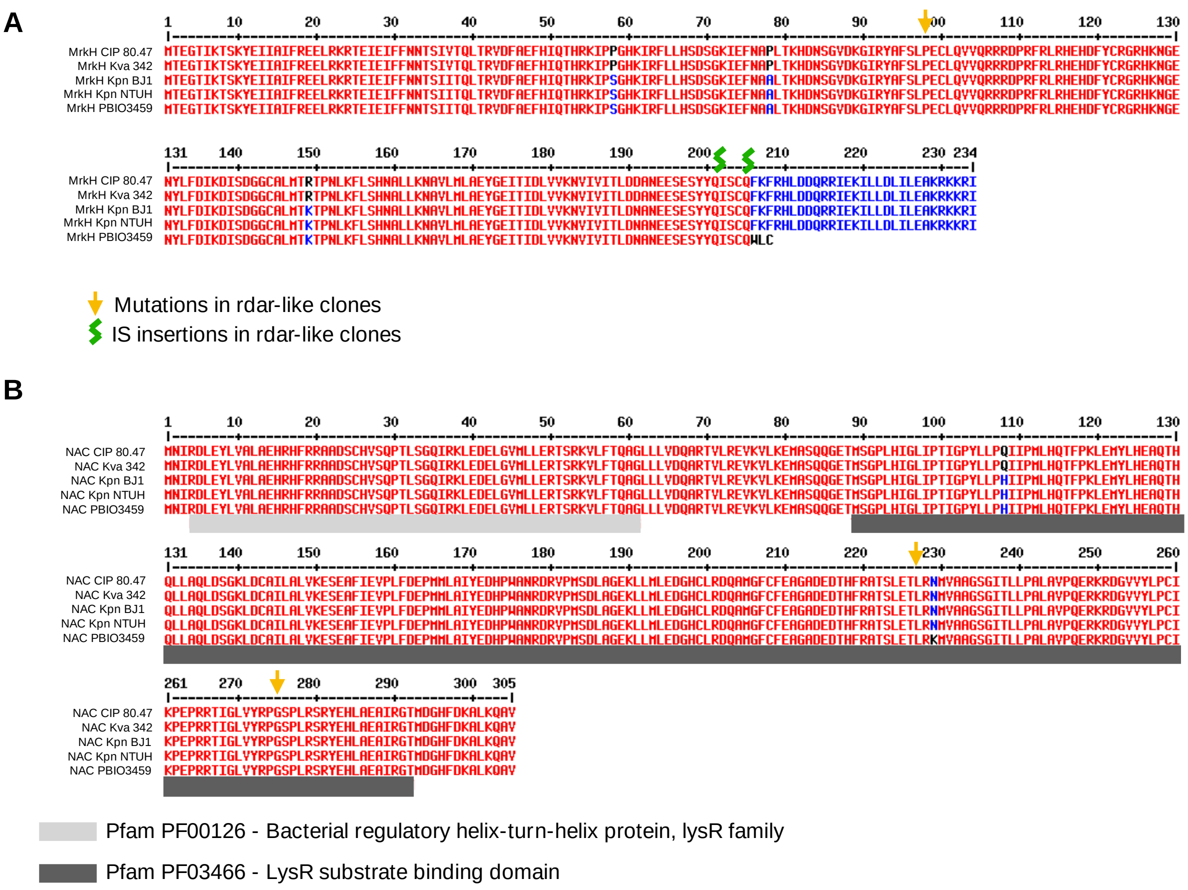


**Figure S7. Alignment of *K. variicola* and *K. pneumoniae* MrkH (A) and NAC (B) proteic sequence.** MrkH and NAC sequences of Kva 342 and PBIO3459 were compared. Two other *K. pneumoniae* (Kpn BJ1 and NTUH) as well as another *K. variicola* CIP 80.47 were included as controls. Graphics and alignments were produced by http://multalin.toulouse.inra.fr/multalin/. Yellow arrows indicate SNPs in our evolution experiment and Green breaks represent insertion of an IS in evolved clones.

### SUPPLEMENTARY TABLES

**Table S1. Accession numbers of protein used as reference to search for the cellulose and curli biosynthesis operon in Kva 342.**

| | Protein | Annotation | Accession |
|---|---|---|---|
| Cellulose biosynthesis operon | BcsA | cellulose synthase catalytic subunit | QJZ01585.1 |
| | BcsB | cellulose biosynthesis cyclic di-GMP-binding regulatory protein | QJZ01584.1 |
| | BcsC | cellulose biosynthesis protein | QJZ01582.1 |
| | BcsE | cellulose biosynthesis protein | QJZ01588.1 |
| | BcsF | | QKU51560.1 |
| | BcsG | | CAD5993002.1 |
| | BcsR | cellulose biosynthesis protein | QJZ01587.1 |
| | BcsQ | cellulose biosynthesis protein | QJZ01586.1 |
| | BcsZ | cellulase | CAD5999681.1 |
| Curli biosynthesis operon | CsgA | major curlin subunit | ACB59307.1 |
| | CsgB | minor curlin subunit | ACB59306.1 |
| | CsgC | Curli assembly protein | KAB2536785.1 |
| | CsgD | Transcriptional regulator, LuxR family transcriptional regulator similar to HTH-type | ACB59305.1 |
| | CsgE | curli production assembly/transport protein | ACB59304.1 |
| | CsgF | curli production assembly/transport protein | ACB59303.1 |
| | CsgG | curli production assembly/transport protein | ACB59302.1 |

**Table S2. Mutations identified in all sequenced clones.** Differences between evolved and ancestral genomes were identified using *breseq* v 0.30.0 (1) with default options.

| Clone | Environment | Replicon | Position | Mutation | Annotation | Gene | Description |
|---|---|---|---|---|---|---|---|
| 4D6 | M02 | Chromosome | 844,705 | IS | | *mrkH* | |
| 4D6 | M02 | Chromosome | 1,448,843 | G→A | R218R (CGC→CGT) | *mnmC* ← | tRNA 5-methylaminomethyl-2-thiouridine biosynthesis bifunctional protein MnmC |
| 4D6 | M02 | Chromosome | 1,847,127 | IS | | *CDS_01717* | hypothetical protein |
| 4D6 | M02 | Chromosome | 4,086,144 | A→T | F93Y (TTC→TAC) | *ramA* ← | |
| 4D6 | M02 | Chromosome | 5,362,500 | C→A | A88D (GCC→GAC) | *thiG* → | Thiazole synthase |
| 6B1 | AUM | Chromosome | 5,497,313 | C→T | G275S (GGC→AGC) | *nac*← | Nitrogen assimilation control protein |
| 6B3 | AUM | Chromosome | 4086431 | IS | | *ramA* | |
| 6B3 | AUM | Chromosome | 687,757 | +T | coding (406/828 nt) | *cpdA_1* → | 3',5'-cyclic adenosine monophosphate phosphodiesterase CpdA |
| 6B3 | AUM | Chromosome | 845,042 | G→A | P98S (CCT→TCT) | *mrkH* ← | |
| 6B3 | AUM | Chromosome | 5,081,979 | G→T | intergenic (-86/+204) | *CDS_04817* ← / ← *cybC* | hypothetical protein/Soluble cytochrome b562 |
| 6B3 | AUM | Plasmid | 49,840 | G→A | E349K (GAA→AAA) | *CDS_05399* → | hypothetical protein |
| 6B4 | AUM | Chromosome | 525,105 | T→C | V326A (GTC→GCC) | *arcB* → | Aerobic respiration control sensor protein ArcB |
| 6B4 | AUM | Chromosome | 1,461,357 | G→A | D52N (GAT→AAT) | *CDS_01391* → | putative membrane protein |
| 6B4 | AUM | Chromosome | 1,983,229 | G→A | intergenic (-40/-298) | *hexR_1* ← / → *zwf* | HTH-type transcriptional regulator HexR/Glucose-6-phosphate 1-dehydrogenase |
| 6B4 | AUM | Chromosome | 4,086,303 | +T | coding (119/342 nt) | *ramA* ← | |
| 6B4 | AUM | Chromosome | 5,498,183 | C→A | intergenic (-48/+212) | *nac* ← / ← *argH* | Nitrogen assimilation control protein /Argininosuccinate lyase |

**Table S3: Frequency table of protein length of MrkH.** MrkH homologs were identified with blastP. Results with an identity of more than 80% were kept and an e-value of <10e-$^{5}$. Wildtype MrkH is 234 amino acids.

| | *K. variicola* | | *K. pneumoniae* | |
|---|---|---|---|---|
| **# of Aminoacids** | **Frequency** | **Prevalence** | **Frequency** | **Prevalence** |
| 0 - 25 | 0 | 0.00 | 0 | 0.00 |
| 25 - 50 | 0 | 0.00 | 3 | 0.03 |
| 50 - 75 | 2 | 0.29 | 42 | 0.44 |
| 75 - 100 | 5 | 0.73 | 663 | 6.96 |
| 100 - 125 | 4 | 0.59 | 33 | 0.35 |
| 125 - 150 | 0 | 0.00 | 8 | 0.08 |
| 150 - 175 | 1 | 0.15 | 23 | 0.24 |
| 175 - 200 | 0 | 0.00 | 31 | 0.33 |
| 200 - 233 | 1 | 0.15 | 64 | 0.67 |
| 233 - 234 | 656 | 96.33 | 7916 | 83.05 |
| 234 - 250 | 0 | 0.00 | 4 | 0.04 |
| MrkH Border | 12 | 1.76 | 839 | 7.81 |
| **Total sequences** | **676** | | **9532** | |

**Table S4. Details of IS elements found in *mrkH* genes.** IS elements were identified using ISFinder (2). Only hits with an e-value of <10e-$^{5}$ were considered. For each IS found overlapping coordinates (including +100 or -100 bp) with those of MrkH, only the best hit per genome was taken into account.

| **Strain** | **PATRIC ID** | **Isolation** | **Source** | **Geographic location** | **IS family** | ***mrkH* coordinates** | **IS coordinates** |
|---|---|---|---|---|---|---|---|
| Kpn 1_19 | 573.30807 | Urine | Human | St. Louis Missouri, USA | IS102_IS5_IS903 | Contig 23:116892-117371 | Contig 23:117341-117417 |
| Kpn 10 | 573.12961 | | Human | Houston, TX, USA | ISEcl1_IS3_IS2 | Contig 32:59770-60492 | Contig 32:60416-60492 |
| Kpn 1164/2047 | 72407.913 | Urine | Human | Michigan, USA | IS1F_IS1_unknown | Contig 14:375744-376439 | Contig 14:376359-377126 |
| Kpn 1247/2119 | 72407.916 | Urine | Human | Michigan, USA | IS1F_IS1_unknown | Contig 27:369766-370461 | Contig 27:370381-371148 |
| Kpn 1281/2159 | 72407.919 | Respiratory | Human | Michigan, USA | IS1F_IS1_unknown | Contig 32:375744-376439 | Contig 32:376359-377126 |
| Kpn 1298/2179 | 72407.969 | Urine | Human | Michigan, USA | IS1F_IS1_unknown | Contig 29:375744-376439 | Contig 29:376359-377126 |
| Kpn 1394/2287 | 72407.1009 | wound | Human | Ohio, USA | ISEc21_IS110_unknown | Contig 38:249417-249686 | Contig 38:249624-250997 |
| Kpn 1382/2273 | 72407.1054 | wound | Human | Ohio, USA | ISEc21_IS110_unknown | Contig 51:249419-249688 | Contig 51:249626-250999 |
| Kpn 1433/2338 | 72407.956 | Respiratory | Human | Michigan, USA | IS1F_IS1_unknown | Contig 10:375744-376439 | Contig 10:376359-377126 |
| Kpn 1463/2375 | 72407.942 | Urine | Human | Michigan, USA | IS1F_IS1_unknown | Contig 81:146931-147626 | Contig 81:146244-147011 |
| Kpn 1472/2387 | 72407.1002 | Urine | Human | Michigan, USA | IS1F_IS1_unknown | Contig 12:375744-376439 | Contig 12:376359-377126 |
| Kpn 150 (ST45) | 573.32426 | | Human | Paris, France | IS903B_IS5_IS903 | Contig 29:63-233 | Contig 29:1-77 |
| Kpn 1533/2460 | 72407.920 | | Human | | IS1F_IS1_unknown | Contig 12:136622-137263 | Contig 12:135935-136702 |
| Kpn 1584/2521 | 72407.96 | Urine | Human | Ohio, USA | ISKpn14_IS1_unknown | Contig 22:84746-85135 | Contig 22:85131-85230 |
| Kpn 1606/2552 | 72407.963 | Urine | Human | Ohio, USA | ISKpn14_IS1_unknown | Contig 28:85387-85776 | Contig 28:85772-85871 |
| Kpn 1851/2844 | 72407.968 | Respiratory | Human | Michigan, USA | IS1F_IS1_unknown | Contig 08:375744-376439 | Contig 08:376359-377126 |
| Kpn 1983/2989 | 72407.971 | Wound | Human | Michigan, USA | IS1F_IS1_unknown | Contig 09:369993-370688 | Contig 09:370608-371375 |

| | | | | | | | |
|---|---|---|---|---|---|---|---|
| Kpn 1983/3063 | 72407.945 | Urine | Human | Michigan,USA | IS1F_IS1_unknown | Contig 93:313881-314576 | Contig 93:314496-315263 |
| Kpn 2000/3010,1 | 72407.943 | Urine | Human | Michigan, USA | IS1F_IS1_unknown | Contig 25:369766-370461 | Contig 25:370381-371148 |
| Kpn 2000/3010,2 | 72407.970 | Urine | Human | Michigan,USA | IS1F_IS1_unknown | Contig 77:369993-370688 | Contig 77:370608-371375 |
| Kpn 20071685 | 573.35693 | | Human | Jiaozou, China | ISKpn26_IS5_IS5 | Contig 34:113-523 | Contig 34:1-127 |
| Kpn 2102/3132 | 72407.974 | Respiratory | Human | Michigan, USA | IS1F_IS1_unknown | Contig 51:122606-123301 | Contig 51:123221-123988 |
| Kpn 2143/3182 | 72407.973 | Urine | Human | Michigan, USA | IS1F_IS1_unknown | Contig 31:369993-370688 | Contig 31:370608-371375 |
| Kpn 2511 | 573.30980 | Conjunctiva | Human | Saint Petesburg, Russia | IS1X2_IS1_unknown | Contig 02:93-377 | Contig 02:1-97 |
| Kpn 2512 | 573.30988 | Conjunctiva | Human | Saint Petesburg, Russia | IS1R_IS1_unknown | Contig 01:258187-258471 | Contig 01:257424-258191 |
| Kpn 285 | 573.24275 | Urine | Human | Boston,MA, USA | IS1S_IS1_unknown | Contig 05:248845-249276 | Contig 05:249196-249322 |
| Kpn 3263 | 573.12905 | | Human | Houston, TX, USA | ISKpn26_IS5_IS5 | Contig 19:119576-120043 | Contig 19:119972-120048 |
| Kpn 39885 | 573.12556 | Blood | Human | USA | ISKpn26_IS5_IS5 | Contig 80:55-561 | Contig 80:1-77 |
| Kpn 4300STDY6470478 | 573.20352 | | Human | Thailand | ISKpn14_IS1_unknown | Contig 05:167-811 | Contig 05:1-125 |
| Kpn 4300STDY6636943 | 573.20205 | | Human | Thailand | ISEc21_IS110_unknown | Contig 06:59535-59804 | Contig 06:58224-59597 |
| Kpn 44810 | 573.12539 | Blood | Human | USA | ISKpn14_IS1_unknown | Contig 18:249762-250262 | Contig 18:250258-250334 |
| Kpn 59 | 573.14774 | Purulent material | Human | Italy | ISKpn26_IS5_IS5 | Contig 77:1150-1875 | Contig 77:1-1196 |
| Kpn 6-83 | 573.35578 | Intra-abdominal infection | Human | Porto, Portugal | ISKpn26_IS5_IS5 | Contig 15:2-655 | Contig 15:1-77 |
| Kpn AS012306 | 573.31220 | Lung | Human | USA | IS903B_IS5_IS903 | Contig 30:116850-117212 | Contig 30:117202-117236 |
| Kpn AS012307 | 573.30859 | Lung | Human | USA | IS903B_IS5_IS903 | Contig 32:116850-117212 | Contig 32:117202-117236 |
| Kpn BA34974 | 573.17828 | Blood | Human | Vellore, India | IS1X2_IS1_unknown | Contig 20:100-534 | Contig 20:1-126 |
| Kpn BIDMC 13 | 1328409.3 | Blood | Human | Boston, USA | IS1294_IS91_unknown | Contig 05:458-1171 | Contig 05:1-488 |

| | | | | | | | |
|---|---|---|---|---|---|---|---|
| Kpn blood_2014 | 573.30806 | Blood | Human | St. Louis ,Missouri, USA | IS102_IS5_IS903 | Contig 25:116892-117371 | Contig 25:117341-117417 |
| Kpn C1-02 | 573.31708 | Sputum | Human | Colombia | ISEc21_IS110_unknown | Contig 13:97135-97404 | Contig 13:97342-98715 |
| Kpn CFSAN059625 | 573.34022 | Urine | Human | Karachi, Pakistan | ISKpn14_IS1_unknown | Contig 11:200308-200796 | Contig 11:200716-200842 |
| Kpn CRE18 | 573.19962 | Blood | Human | Palo Alto, CA, USA | IS1F_IS1_unknown | Contig 02:183645-183941 | Contig 02:182899-183666 |
| Kpn CriePir134 | 573.28668 | Blood | Human | Moscow, Russia | IS1R_IS1_unknown | Contig 03:133498-134142 | Contig 03:134201-134968 |
| Kpn CriePir183 | 573.28655 | Urine | Human | Moscow, Russia | ISVsa5_IS4_IS10 | Contig 14:121-813 | Contig 14:1-124 |
| Kpn CriePir187 | 573.28662 | Traumatic discharge | Human | Moscow, Russia | IS1R_IS1_unknown | Contig 17:186-830 | Contig 17:1-127 |
| Kpn CriePir226 | 573.28649 | Blood | Human | Moscow,Russia | IS1S_IS1_unknown | Contig 05:106-714 | Contig 05:1-127 |
| Kpn CRK0030 | 573.22495 | Blood | Human | North Carolina, USA | IS1F_IS1_unknown | Contig 56:375744-376439 | Contig 56:376359-377126 |
| Kpn CRK0036 | 573.22493 | Blood | Human | Ohio, USA | IS1F_IS1_unknown | Contig 05:375744-376439 | Contig 05:376359-377126 |
| Kpn CRK0074 | 573.22535 | Blood | Human | Ohio, USA | IS1F_IS1_unknown | Contig 60:375744-376439 | Contig 60:376359-377126 |
| Kpn CRK0127 | 573.22455 | Respiratory | Human | Michigan, USA | IS1F_IS1_unknown | Contig 03:319511-320206 | Contig 03:320126-320893 |
| Kpn CRK0131 | 573.22376 | Respiratory | Human | Ohio, USA | ISEc21_IS110_unknown | Contig 30:249644-249913 | Contig 30:249851-251224 |
| Kpn CRK0137 | 573.22382 | Urine | Human | Michigan, USA | IS1F_IS1_unknown | Contig 63:369993-370688 | Contig 63:370608-371375 |
| Kpn CRK0138 | 573.22399 | Urine | Human | Michigan, USA | IS1F_IS1_unknown | Contig 09:375744-376439 | Contig 09:376359-377126 |
| Kpn CRK0152 | 573.22384 | Urine | Human | Michigan, USA | IS1F_IS1_unknown | Contig 21:146931-147626 | Contig 21:146244-147011 |
| Kpn CRK0159 | 573.22370 | Urine | Human | Michigan, USA | IS1F_IS1_unknown | Contig 88:375744-376439 | Contig 88:376359-377126 |
| Kpn CRK0166 | 573.22431 | Urine | Human | Michigan, USA | IS1X2_IS1_unknown | Contig 93:375744-376439 | Contig 93:376359-376496 |
| Kpn CRKP380 | 573.29492 | Blood septicemia | Human | Hangzhou, China | ISKpn14_IS1_unknown | Contig 03:117-461 | Contig 03:1-121 |
| Kpn DJJYY72-CR | 573.26001 | | Human | Jiangsu, China | ISKpn26_IS5_IS5 | Contig 02:475-597 | Contig 02:1-502 |
| Kpn DLL7528 | 573.19849 | Stool | Human | Perth, Australia | IS903B_IS5_IS903 | Contig 20:97184-97354 | Contig 20:97340-97586 |
| Kpn e1602 | 573.4176 | Blood | Human | United Kingdom | IS903B_IS5_IS903 | Contig 15:86-571 | Contig 15:1-100 |

| | | | | | | | |
|---|---|---|---|---|---|---|---|
| Kpn EuSCAPE_FR021 | 1463165.89 | Blood infection | Human | France | IS1R_IS1_unknown | Contig 08:216-335 | Contig 08:1-220 |
| Kpn EuSCAPE_GR073 | 573.20467 | Wound | Human | Greece | IS1S_IS1_unknown | Contig 07:89-400 | Contig 07:1-93 |
| Kpn EuSCAPE_GR075 | 573.20468 | Blood infection | Human | Grece | IS1S_IS1_unknown | Contig 07:89-400 | Contig 07:1-93 |
| Kpn EuSCAPE_H2092 | 573.20876 | Urine | Human | Croatia | IS1S_IS1_unknown | Contig 08:200274-200582 | Contig 08:200561-200653 |
| Kpn EuSCAPE_HR068 | 573.20989 | Low Respiratory tract secretion | Human | Croatia | IS1S_IS1_unknown | Contig 09:72-380 | Contig 09:1-93 |
| Kpn EuSCAPE_HR080 | 573.21001 | Blood | Human | Croatia | IS903B_IS5_IS903 | Contig 10:93-455 | Contig 10:1-93 |
| Kpn EuSCAPE_IT221 | 573.21751 | Blood infection | Human | Italy | IS1R_IS1_unknown | Contig 04:377714-378082 | Contig 04:378002-378769 |
| Kpn EuSCAPE_RO062 | 573.20792 | Puncture fluids | Human | Romania | IS1X2_IS1_unknown | Contig 13:97420-97884 | Contig 13:97880-97972 |
| Kpn EuSCAPE_TR121 | 573.21111 | Blood | Human | Turkey | IS1S_IS1_unknown | Contig 06:89-496 | Contig 06:1-93 |
| Kpn EuSCAPE_TR275 | 573.21296 | Low Respiratory tract secretion | Human | Turkey | IS1X2_IS1_unknown | Contig 09:67-630 | Contig 09:1-93 |
| Kpn GM4 | 573.27962 | Sputum | Human | Shenzhen, China | IS102_IS5_IS903 | Contig 02:290796-291296 | Contig 02:291266-291561 |
| Kpn JS86C16 | 573.1679 | Urine | Human | San Francisco, USA | IS903B_IS5_IS903 | Contig 04:117-698 | Contig 04:1-127 |
| Kpn k1337 | 573.7056 | Blood | Human | United Kingdom | IS903B_IS5_IS903 | Contig 03:127623-127793 | Contig 03:127779-127888 |
| Kpn k1580 | 573.7077 | Blood | Human | United Kingdom | ISEcp1_IS1380_unknown | Contig 77:66-428 | Contig 77:509-584 |

| | | | | | | | |
|---|---|---|---|---|---|---|---|
| Kpn k2 | 573.35638 | Blood | Human | Egypt | IS1S_IS1_unknown | Contig 32:185-829 | Contig 32:1-126 |
| Kpn k2370 | 573.414 | Blood | Human | United Kingdom | IS903B_IS5_IS903 | Contig 02:179-349 | Contig 02:1-193 |
| Kpn K32 | 573.34089 | Blood | Human | South Africa | IS1S_IS1_unknown | Contig 16:141008-141445 | Contig 16:141424-141525 |
| Kpn k589 | 573.4081 | Blood | Human | United Kingdom | IS903B_IS5_IS903 | Contig 54:32961-33131 | Contig 54:33117-33309 |
| Kpn k722 | 573.4183 | Blood | Human | United Kingdom | IS903B_IS5_IS903 | Contig 12:398199-398699 | Contig 12:398669-398895 |
| Kpn k866 | 573.4083 | Blood | Human | United Kingdom | ISCro1_IS66_unknown | Contig 14:51-758 | Contig 14:1-77 |
| Kpn KPM_43 | 573.30889 | | Human | Lebanon | IS1R_IS1_unknown | Contig 09:200308-200871 | Contig 09:200845-200971 |
| Kpn KPN 974ec | 573.14534 | Urine | Human | Houston, TX, USA | IS903B_IS5_IS903 | Contig 23:3-635 | Contig 23:1-77 |
| Kpn KPN1105ec | 573.13109 | Blood | Human | Houston, TX, USA | ISKpn14_IS1_unknown | Contig 08:206378-206686 | Contig 08:206665-206741 |
| Kpn KPN1107ec | 573.13111 | Blood | Human | Houston, TX, USA | ISKpn14_IS1_unknown | Contig 02:56-364 | Contig 02:1-77 |
| Kpn KPN1251ec | 573.13231 | Urine | Human | Houston, TX, USA | ISKpn26_IS5_IS5 | Contig 18:55-756 | Contig 18:1-77 |
| Kpn KPN1265ec | 573.13244 | Lung | Human | Houston, TX, USA | IS903B_IS5_IS903 | Contig 46:249531-250040 | Contig 46:250010-250086 |
| Kpn KPN1378ec | 573.13329 | Blood | Human | Houston, TX, USA | IS903B_IS5_IS903 | Contig 23:3-635 | Contig 23:1-77 |
| Kpn KPN1380ec | 573.13330 | Blood | Human | Houston, TX, USA | IS903B_IS5_IS903 | Contig 23:3-635 | Contig 23:1-77 |
| Kpn KPN1936ec | 573.13787 | Tissue | Human | Houston, TX, USA | ISKpn26_IS5_IS5 | Contig 25:96-584 | Contig 25:1-123 |
| Kpn KPN1994ec | 573.13827 | Respiratory | Human | Houston, TX, USA | IS903B_IS5_IS903 | Contig 63:200039-200401 | Contig 63:200391-200467 |
| Kpn KPN1997ec | 573.13828 | Respiratory | Human | Houston, TX, USA | IS903B_IS5_IS903 | Contig 50:200039-200401 | Contig 50:200391-200520 |
| Kpn KPN2043ec | 573.13869 | Respiratory | Human | Houston, TX, USA | IS903B_IS5_IS903 | Contig 09:200039-200401 | Contig 09:200391-200467 |
| Kpn KPN2101ec | 573.13922 | Urine | Human | Houston, TX, USA | ISKpn26_IS5_IS5 | Contig 16:119659-120384 | Contig 16:120338-120414 |
| Kpn KPN708ec | 573.14335 | Respiratory | Human | Houston, TX, USA | IS903B_IS5_IS903 | Contig 25:67-429 | Contig 25:1-77 |
| Kpn KPN719ec | 573.14345 | Urine | Human | Houston, TX, USA | IS903B_IS5_IS903 | Contig 08:67-429 | Contig 08:1-77 |
| Kpn KPN762ec | 573.14379 | Respiratory | Human | Houston, TX, USA | IS903B_IS5_IS903 | Contig 07:67-429 | Contig 07:1-77 |
| Kpn KPN801ec | 573.14405 | Respiratory | Human | Houston, TX, USA | IS903B_IS5_IS903 | Contig 35:67-429 | Contig 35:1-77 |

| | | | | | | | |
|---|---|---|---|---|---|---|---|
| Kpn KPN811ec | 573.14413 | Urine | Human | Houston, TX, USA | ISKpn26_IS5_IS5 | Contig 37:5-730 | Contig 37:1-51 |
| Kpn KPN814ec | 573.14415 | Respiratory | Human | Houston, TX, USA | IS903B_IS5_IS903 | Contig 07:67-429 | Contig 07:1-77 |
| Kpn KPN816ec | 573.14416 | Respiratory | Human | Houston, TX, USA | IS903B_IS5_IS903 | Contig 06:67-429 | Contig 06:1-77 |
| Kpn KPN913ec | 573.14487 | Ulceer | Human | Houston, TX, USA | ISKpn26_IS5_IS5 | Contig 06:31-756 | Contig 06:1-77 |
| Kpn KPN914ec | 573.14488 | Urine | Human | Houston, TX, USA | ISKpn26_IS5_IS5 | Contig 16:31-756 | Contig 16:1-77 |
| Kpn KPN915ec | 573.14489 | Urine | Human | Houston, TX, USA | ISKpn26_IS5_IS5 | Contig 18:31-756 | Contig 18:1-77 |
| Kpn KPN965ec | 573.14529 | Abscess | Human | Houston, TX, USA | ISVsa5_IS4_IS10 | Contig 30:3-512 | Contig 30:1-77 |
| Kpn KPN980ec | 573.14538 | Urine | Human | Houston, TX, USA | ISVsa5_IS4_IS10 | Contig 36:37212-37721 | Contig 36:37647-37723 |
| Kpn L61 | 573.14880 | Stool | Human | Zhejiang, China | ISKpn26_IS5_IS5 | Contig 03:304277-304924 | Contig 03:304897-305023 |
| Kpn MGH 80 | 1438801.3 | Urine | Human | Boston, MA, USA | IS1X2_IS1_unknown | Contig 13:98066-98710 | Contig 13:97240-98007 |
| Kpn MGH171 | 573.9832 | | Human | Boston, MA, USA | IS1R_IS1_unknown | Contig 09:62706-63269 | Contig 09:63243-64010 |
| Kpn MUGSI_242 | 573.24354 | Blood | Human | USA | IS903_IS5_IS903 | Contig 14:154431-154886 | Contig 14:154872-155171 |
| Kpn NMI759_17 | 573.31649 | Throat | Human | Poland | IS903B_IS5_IS903 | Contig 52:416-676 | Contig 52:666-790 |
| Kpn NNKP315 | 573.32126 | Wound | Human | Nizhniy Novgorod, Russia | IS1S_IS1_unknown | Contig 20:2-514 | Contig 20:1-55 |
| Kpn NNKP343 | 573.32127 | Wound | Human | Nizhniy Novgorod, Russia | IS1X2_IS1_unknown | Contig 79:113-757 | Contig 79:1-54 |
| Kpn NR5099 | 573.3329 | | Human | New York, USA | ISKpn14_IS1_unknown | Contig 01:446869-447369 | Contig 01:446106-446873 |
| Kpn NR5391 | 573.33292 | | Human | New York, USA | ISKpn14_IS1_unknown | Contig 01:339540-340040 | Contig 01:340036-340803 |
| Kpn NR5476 | 573.33352 | | Human | New York, USA | ISKpn14_IS1_unknown | Contig 01:339540-340040 | Contig 01:340036-340803 |
| Kpn SCKP020082 | 573.18225 | | Human | Sichuan, China | IS903B_IS5_IS903 | Contig 11:89-493 | Contig 11:1-99 |
| Kpn SHK038 | 573.27562 | secretion | Human | Shangai, China | ISKpn14_IS1_unknown | Contig 12:123-608 | Contig 12:1-127 |
| Kpn SHK072 | 573.27555 | Sputum | Human | Shangai, China | ISKpn14_IS1_unknown | Contig 13:200308-200793 | Contig 13:200789-200915 |
| Kpn SHK128 | 573.27557 | Secretion | Human | Shangai, China | ISKpn14_IS1_unknown | Contig 10:123-608 | Contig 10:1-127 |

| | | | | | | | |
|---|---|---|---|---|---|---|---|
| Kpn SIKP009 | 72407.1300 | Stool | Human | Thailand | ISEcp1_IS1380_unknown | Contig 22:37-639 | Contig 22:1-77 |
| Kpn SIKP107 | 72407.1360 | Stool | Human | Thailand | ISEc68_IS5_IS5 | Contig 17:46-651 | Contig 17:1-73 |
| Kpn ST258 | 573.1757 | Urine | Human | Brisbane, Australia | ISEc21_IS110_unknown | Contig 10:97128-97397 | Contig 10:97335-98708 |
| Kpn TX001-S00-A | 573.30811 | Stool | Human | Missouri, USA | IS102_IS5_IS903 | Contig 41:47-526 | Contig 41:1-77 |
| Kpn TX001-S01 | 573.30808 | Stool | Human | St. Louis, Missouri,USA | IS102_IS5_IS903 | Contig 25:116895-117374 | Contig 25:117344-118010 |
| Kpn TX001-S05 | 573.30810 | Stool | Human | St. Louis, Missouri,USA | IS102_IS5_IS903 | Contig 22:47-526 | Contig 22:1-77 |
| Kpn urine_4_26 | 573.30805 | Urine | Human | St. Louis, Missouri,USA | IS102_IS5_IS903 | Contig 50:47-526 | Contig 50:1-77 |
| Kpn WCHKP85522 | 573.14848 | | Human | Sichuan, China | IS5D_IS5_IS5 | Contig 05:344657-345145 | Contig 05:345118-345191 |
| Kpn WUSM_KP_113 | 573.18491 | Blood | Human | St. Louis, Missouri,USA | IS903_IS5_IS903 | Contig 05:476-646 | Contig 05:4-490 |
| Kpn WUSM_Kp_65 | 573.18515 | Urine | Human | St. Louis, Missouri,USA | IS102_1S5_IS903 | Contig 52: 20691-21170 | Contig 52: 21140-21216 |
| Kpn ZS15 | 573.32390 | Urine | Human | | ISKpn26_IS5_IS5 | Contig 54:94-699 | Contig 54:1-121 |
| Kvar 8917 | 244366.23 | Expectoration | Human | Mexico | IS903_IS5_IS903 | Contig 5: 476-646 | Contig 5: 4-490 |
| Kvar KP-21F | 244366.89 | Blood | Human | Groningen, Netherlands | IS1R_IS1_unknown | Contig 52: 747-1274 | Contig 52: 1-768 |
| K. quasipneumoniae B8095 | 1463165.19 | Blood bacteriemia | Human | India | IS102_IS5_IS903 | Contig 08:89-451 | Contig 08:1-99 |
| K. quasipneumoniae C1-124 | 1463165.26 | Rectal swab | Human | Colombia | IS903B_IS5_IS903 | Contig 94:34231-34437 | Contig 94:34423-34615 |
| K. quasipneumoniae TUM14048 | 1463165.192. | Blood | Human | Japan | IS903B_IS5_IS903 | Contig 05:218436-218555 | Contig 05:218541-218665 |

#### Table S5. Primers used in this study.

| # | Name | Sequence | Direction | Comments |
|---|---|---|---|---|
| qPCR primers | | | | |
| 657 | recA.qPCR_new.5 | CATCAACTTCTTCGGCGAGC | Forward | *recA* amplification for qPCR |
| 357 | recA.qPCR.new.3 | CGCGAACTTTCTTCTCAATTTCT | Reverse | *recA* amplification for qPCR |
| 320 | rho.qPCR.5 | GGAGATACTGCAGGATGGATTT | Forward | *rho* amplification for qPCR |
| 321 | rho.qPCR.3 | GGTTGAAACGGCGGATTTG | Reverse | *rho* amplification for qPCR |
| 633 | mrkA127F | AGCGATGCGAACGTTTACCTGTCTC | Forward | *mrkA* amplification for qPCR |
| 634 | mrkA265R | cgtcatcctgtttggtgccgtcagc | Reverse | *mrkA* amplification for qPCR |
| Mutant construction | | | | |
| 613 | 24_500_mrkH_F | CATAAGTAGAAGCAGCAACCCAAGTAG CTTTACCAGCATCcctctataacattgcccatcgc | Forward | *mrkH* deletion or mutation in Kva 342 + pKNG101 tail |
| 614 | 24_500_mrkH_R | CTTCCGCTCAGGTCCTTGTCCTTTAACG AGGATTGTTACaggttagtcagacaataggc | Reverse | *mrkH* deletion or mutation in Kva 342 + pKNG101 tail |
| 615 | 24_mrkH_atg_R | gtgcttcccttgtaaatagttgtcg | Reverse | *mrkH* deletion in Kva 342 |
| 616 | 24_mrkH_stop_F | AAATCAAACGCCTCACGACAACTATTTA CAAGGGAAGCACtacccttggacagtatattgctg | Forward | *mrkH* deletion in Kva 342 |
| 617 | 24_mrkH_out_F | caggtttatcggttgggttatgg | Forward | Deletion verification in *mrkH* |
| 618 | 24_mrkH_out_R | ccctctgatggcgttaatcc | Reverse | Deletion verification in *mrkH* |
| 660 | Nac_Gibson5 | CATAAGTAGAAGCAGCAACCCAAGTAG CTTTACCAGCATCtcacaggtttaccgtctctatgg | Forward | Amplification of *nac* + pKNG101 tail |
| 661 | Nac_Gibson3 | CTTCCGCTCAGGTCCTTGTCCTTTAACG AGGATTGTTACcaccacctttaactacccgacc | Reverse | Amplification of *nac* + pKNG101 tail |
| 438 | pkng101.R gibson | GTAACAATCCTCGTTAAAGGAC | Reverse | linearize pKNG101 for gibson |
| 439 | pkng101.F gibson | GATGCTGGTAAAGCTACTTG | Forward | linearize pKNG101 for gibson |
| Sequence verification | | | | |
| 629 | mrkH_Sanger_F | AACCCAATCAGAATGTTGC | Forward | *mrkH* Sanger sequencing |
| 630 | mrkH_Sanger_R | GTAGTGATAGATTGAGTGACC | Reverse | *mrkH* Sanger sequencing |
| 658 | Nac_verif5 | GTTCACGTTGGCTTTAGTTGTTCG | Forward | *nac* Sanger sequencing |
| 659 | Nac_verif3 | ACCCTGAAGATCCTGTTCCATCG | Reverse | *nac* Sanger sequencing |
| 532 | ramA_verif5 | ATGAAACGGCTCAGGCTGC | Forward | *ramA* Sanger sequencing |
| 533 | ramA_verif3 | CTCTCCTGTCTGCTGTTGC | Reverse | *ramA* Sanger sequencing |
